# Transcriptome analysis reveals role for WRKY70 in early *N-*hydroxy-pipecolic acid signaling

**DOI:** 10.1101/2024.04.23.590810

**Authors:** Jessica Foret, Jung-Gun Kim, Elizabeth S. Sattely, Mary Beth Mudgett

## Abstract

*N*-hydroxy-pipecolic acid (NHP) is a mobile metabolite essential for inducing and amplifying systemic acquired resistance (SAR) following pathogen attack. Early phases of NHP signaling leading to immunity have remained elusive. Here we report the early transcriptional changes mediated by NHP and the role salicylic acid (SA) plays during this response. We show that distinct waves of expression within minutes to hours of NHP treatment include increased expression of WRKY transcription factors as the primary transcriptional response, followed by the induction of WRKY-regulated defense genes as the secondary response. The majority of genes induced by NHP within minutes were SA-dependent, whereas those induced within hours were SA-independent. These data suggest that NHP induces the primary transcriptional response in a low SA environment and new SA biosynthesis is dispensable for induction of the secondary transcriptional response. We demonstrate that WRKY70 is required for the induced expression of a set of genes defining some of the secondary transcriptional response, SAR protection, and NHP-dependent enhancement of ROS production in response to flagellin treatment. Taken together, our study highlights the key genes and pathways defining early NHP responses and a role for WRKY70 in the regulation of NHP-dependent transcription.

## Introduction

Systemic acquired resistance (SAR) is a whole-plant defense response triggered by an initial pathogen attack. During the establishment of SAR, mobile signals are sent from infected tissue to uninfected tissue, initiating the priming of immune responses and enhancing immunity against secondary pathogen infections (Fu and Dong, 2013; Tian and Zhang, 2019; Kachroo and Kachroo, 2020). To achieve a primed state, plants execute substantial transcriptional reprogramming of several hundred to a few thousand genes, resulting in the increased accumulation of defense hormone salicylic acid (SA), sensing and signaling components for pathogen detection, and production of antimicrobial compounds (Jaskiewicz et al., 2011; Conrath et al., 2015; Bernsdorff et al., 2016). In addition, several small molecules produced by plants during infection help to orchestrate this defense priming, including SA and the bioactive metabolite *N*-hydroxy-pipecolic acid (NHP) (Chen et al., 2018; Hartmann et al., 2018; Klessig et al., 2018).

NHP is a mobile metabolite that can move from one leaf to another to mediate long-distance cellular communication (Chen et al., 2018; Mohnike et al., 2021; Yildiz et al., 2021). In *Arabidopsis thaliana* (Arabidopsis), the enzyme FLAVIN-DEPENDENT MONOOXYGENASE 1 (FMO1) catalyzes the *N*-hydroxylation of lysine-derived pipecolic acid (Pip) in response to pathogen infection to produce NHP (Chen et al., 2018; Hartmann et al., 2018). Notably, *fmo1* null mutants are impaired in local disease resistance during pathogen infection and fail to establish SAR (Chen et al., 2018; Hartmann et al., 2018). Treatment of *fmo1* mutant plants with exogenous NHP is sufficient to induce systemic resistance against bacterial and oomycete pathogens (Chen et al., 2018; Hartmann et al., 2018). Furthermore, NHP treatment of a single leaf is sufficient to initiate and amplify defense signaling throughout the plant, resulting in the transcriptional activation of a large subset of SAR-associated genes within 24 h of treatment (Yildiz et al., 2021; Yildiz et al., 2023), including genes encoding the SA biosynthetic enzyme ISOCHORISMATE SYNTHASE 1/SA-INDUCTION DEFICIENT 2 (ICS1/SID2), the Pip biosynthetic enzymes AGD2-LIKE DEFENSE RESPONSE PROTEIN 1 (ALD1) and SAR-DEFICIENT 4 (SARD4), as well as its own biosynthetic enzyme FMO1 (Chen et al., 2018). Collectively, these findings highlight the importance of NHP in initiating immune signaling to establish local and systemic plant defenses.

Induction of *ICS1* transcription by NHP signaling results in the accumulation of SA in both local and systemic tissues (Chen et al., 2018; Hartmann et al., 2018). Elevation of SA in turn amplifies SAR-associated transcriptional changes (Bernsdorff et al., 2016). Signaling downstream of SA accumulation is largely driven by the action of the SA receptor and transcriptional coactivator NONEXPRESSER OF PR GENES 1 (NPR1), which interacts with a number of TGA transcription factors (TFs) from the basic leucine zipper (bZIP) family to alter gene expression (Zhang et al., 1999; Wang et al., 2006; Wu et al., 2012). Members of the biotic and abiotic stress-response WRKY TF family are among the known targets of the NPR1/TGA interaction (Thibaud-Nissen et al., 2006; Wang et al., 2006). WRKY TFs have been identified as key regulators of NPR1-dependent and NPR1-independent SAR responses. Additionally, some WRKY members play important roles in feedback regulation of transcription (Pandey and Somssich, 2009; Fu and Dong, 2013). For example, WRKY54 and WRKY70 modulate SA biosynthesis through repression of *ICS1* expression (Wang et al., 2006).

The interplay between SA and NHP appears to be interdependent and synergistic with both metabolites being able to upregulate one another’s biosynthetic pathways (Chen et al., 2018; Hartmann et al., 2018; Sun et al., 2020). Notably, elevated endogenous levels of Pip, the precursor of NHP, primes the expression of both NHP and SA biosynthetic genes under low concentrations of SA by stabilizing NPR1 levels (Kim et al., 2020). This points to an intriguing dynamic relationship between SA and NHP over the course of SAR activation. Evidence suggests NHP accumulates to detectable levels in systemic tissues roughly 24 h before SA accumulation is detected (Hartmann and Zeier, 2019). In such a case, NHP would need to drive systemic signaling for a substantial period of time before SA could further amplify SAR responses. It is currently unclear how the levels of SA in leaves impact early NHP signaling. Furthermore, little is known about the very early phase of NHP signaling in the tissues where it is first produced or in the distal tissues once NHP arrives there. Key players of NHP signaling and the progression of NHP-dependent transcriptional reprogramming require further investigation.

Presently, the majority of genome-wide transcriptomic studies of SAR have been carried out on the timescale of 1-2 days post induction; this includes pharmacological studies using Pip and NHP treatment (Bernsdorff et al., 2016; Hartmann et al., 2018; Yildiz et al., 2021; Yildiz et al., 2023). These data have revealed that NHP ultimately activates SAR in untreated tissue, but do not shed light on early signaling events that lead to the observed SAR phenotype. The immediate consequences of NHP accumulation on gene expression have not been described. Studying very early transcriptional changes, within minutes to hours of elicitor treatment, has been an effective strategy elucidating key components of signaling pathways for SA, jasmonic acid (JA), and brassinosteroid (Friedrichsen et al., 2002; Chini et al., 2007; Ding et al., 2018). Thus, in an effort to better understand NHP’s primary action in signal transduction, we determined the genome-wide transcriptional responses in Arabidopsis from 15 minutes up to 6 hours after treatment with exogenous NHP.

Here we report the early NHP transcriptional profile and the role SA plays during this response. We present evidence indicating that increased expression of WRKY TFs define the primary transcriptional responses mediated by NHP within minutes (i.e. 15 to 30 minutes) post-treatment, and WRKY TFs act as executors of the secondary transcriptional responses within hours (i.e. 3 to 6 hours). The majority of genes induced by NHP within minutes were SA-dependent, whereas those induced within hours were SA-independent. These data suggest that NHP induces the primary transcriptional response in a low SA environment and new SA biosynthesis is dispensable for induction of the secondary transcriptional response. We also show that WRKY70 is required for NHP induced transcription of a set of genes defining secondary transcriptional changes as well as NHP enhancement of reactive oxygen species (ROS) and SAR. Taken together, our study highlights the key genes and pathways defining early NHP responses and a role for WRKY70 in the regulation of NHP-dependent transcription.

## Results

### NHP induces expression of defense-associated genes within minutes

Previous studies in Arabidopsis have demonstrated that exogenous application of NHP to a leaf (defined as “local tissue”) is sufficient to initiate and amplify defense signaling in untreated leaves (defined as “distal tissue”), including the transcriptional activation of SAR marker genes (Chen et al., 2018; Hartmann et al., 2018; Nair et al., 2021; Yildiz et al., 2021). Notably, these studies focused on gene expression changes in distal tissues at late time points (24 to 48 h) following NHP elicitation. However, it was unknown which gene expression changes occur immediately in response to NHP treatment. We hypothesized that once present in the tissue, either via endogenous synthesis or exogenous application, NHP initiates signaling that leads to transcriptional changes on a timescale of minutes to several hours, which are essential for execution of NHP-dependent immune responses.

In order to test this hypothesis, we set out to identify NHP responsive transcriptional markers and determine the earliest time frame when NHP could induce these genes. We first questioned if NHP could induce the expression of classical SAR-associated marker genes (i.e., *FMO1*, *ICS1*, and *PATHOGENESIS-RELATED PROTEIN 1* (*PR1*)) within several hours of treatment. Studies have shown *FMO1*, *ICS1*, and *PR1* are induced between 8 to 48 h following NHP treatment (Chen et al., 2018; Nair et al., 2021). Therefore, we began by testing the response to NHP within 24 h of treatment. The leaves of 4.5-week-old Arabidopsis Col-0 (wild type) plants were infiltrated with 1 mM NHP and then treated leaves were collected for mRNA isolation at 2, 6 and 24 h. Quantitative real-time PCR (qRT-PCR) analysis showed significantly increased transcript abundance of all three genes in NHP treated leaves at 24 h but not at 2 and 6 h compared to water (mock) treated leaves (Fig. 1A). These data indicate that *FMO1*, *ICS1,* and *PR1* are responsive to NHP treatment but are likely not primary targets of NHP transcriptional activation.

**Figure 1.**
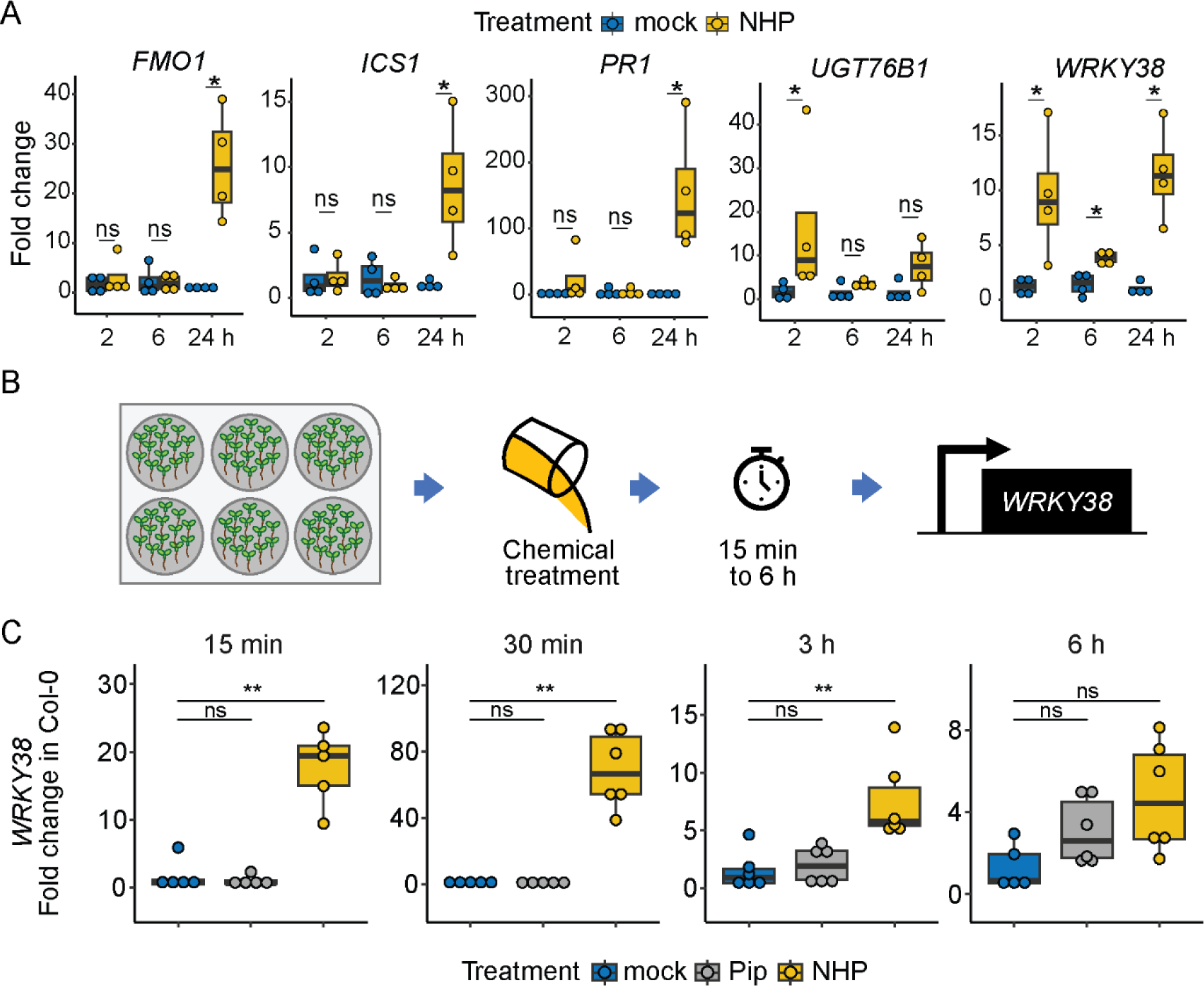
NHP induces *WRKY38* expression at early time points. A, Transcript abundance of SAR marker genes *FMO1*, *ICS1*, and *PR1* and early SA-responsive marker genes *WRKY38* and *UGT76B1* in 4.5-week-old wild type (Col-0) Arabidopsis plants. Three leaves were infiltrated with sterile water (mock) or 1 mM NHP and collected after 2, 6, and 24 h for mRNA isolation. Transcript abundance was measured via quantitative real-time PCR (qRT-PCR) and normalized relative to *UBC21* (Ubiquitin-Conjugating Enzyme 21; At5g25760). Fold change was determined relative to mock (2^−ΔΔCt^). Asterisks indicate a significant difference between mock and NHP per time point (n = 4; Mann-Whitney *U* test; \**P* < 0.05, ns = not significant). B, Design of seedling-based assay. Arabidopsis seedlings (15-18 per well) were hydroponically grown, treated with MS medium (mock), 1 mM Pip, or 1 mM NHP in MS medium then collected at 15 min, 30 min, 3 h, and 6 h for qRT-PCR analysis of *WRKY38* transcript abundance in wild type. C, Expression of *WRKY38* in seedlings from the experiment described in B. Transcript abundance was normalized relative to *UBC21* and mock treated samples for each condition (2^−ΔΔCt^). Asterisks indicate a significant difference between mock and Pip or NHP at each time point (n = 4-6; Mann-Whitney *U* test; \*\**P* < 0.01, ns = not significant).

Next, we tested the ability of NHP to induce expression of *WRKY38* and *UGT76B1*, two genes known to respond to SA within 2 h of treatment (Blanco et al., 2009). *WRKY38* encodes a defense-associated TF and *UGT76B1* encodes a UDP-dependent glycosyltransferase shown to glycosylate SA and NHP (Eulgem et al., 2000; Bauer et al., 2021; Cai et al., 2021; Holmes et al., 2021; Mohnike et al., 2021). We found a significant increase of mRNA abundance for both *WRKY38* and *UGT76B1* in NHP treated leaves at 2 h compared to mock treated leaves (Fig. 1A). *WRKY38* transcript levels were also elevated at 6 and 24 h after treatment with NHP (Fig. 1A). These data indicate that both *WRKY38* and *UGT76B1* are early NHP responsive transcripts.

We used the *WRKY38* gene as a marker to define the earliest time when NHP-induced transcriptional changes could be measured. To this end, we implemented a hydroponically grown seedling assay to synchronize chemical treatment and increase the number of plants sampled, while minimizing mechanical stress. Wild type seedlings were grown in liquid medium in six-well plates with 15-18 seeds per well. The seedlings were treated with mock, 1 mM Pip, or 1 mM NHP at 10-days-post germination and then collected at 15 min, 30 min, 3 h, and 6 h post treatment to isolate mRNA (Fig. 1B). *WRKY38* transcript abundance was significantly elevated in Arabidopsis seedlings treated with NHP as early as 15 min, peaking in abundance at 30 min, when compared to those treated with mock or Pip (Fig. 1C). This trend continued through 3 h following NHP treatment. The NHP precursor Pip did not elicit an increase in *WRKY38* mRNA within 3 h but showed a slight trend of increased abundance at the 6 h time point (Fig. 1C). These results indicate that exogenous NHP, but not Pip, is sufficient to elevate *WRKY38* transcript abundance within 15 minutes of application.

Since *WRKY38* expression is known to be induced by SA treatment (Blanco et al., 2009), we questioned if NHP-induced expression of *WRKY38* is dependent on SA derived from the ICS1/SID2 biosynthetic pathway by utilizing the well characterized *sid2-2* mutant that does not accumulate SA upon pathogen infection (Wildermuth et al., 2001). Wild type and *sid2-2* seedlings were grown hydroponically, treated with mock, 1 mM Pip, or 1 mM NHP, and then collected 3 h after treatment. NHP treatment increased *WRKY38* mRNA abundance in the *sid2-2* mutant relative to mock treatment, similar to wild type seedlings (Supplemental Fig. S1). These findings indicate that NHP is sufficient to alter *WRKY38* transcript abundance in the absence of SA accumulation, defining *WRKY38* as an early NHP induced, SA-independent transcript.

### The early NHP-responsive transcriptome reveals distinct waves of expression

We next questioned how the presence of NHP impacts whole-genome transcriptional changes within minutes to hours of treatment in a SA-dependent and -independent manner. To address this, Arabidopsis wild type and *sid2-2* mutant seedlings were grown in the same hydroponic design as previously described for qRT-PCR experiments (Fig. 1B). The *sid2-2* mutant was analyzed in order to identify transcriptional changes dependent on NHP-induced SA biosynthesis (Chen et al., 2018). Each pool of seedlings was treated with mock or 0.5 mM NHP for 15 min, 30 min, 3 h and 6 h. A reduced concentration of NHP was used as we found that 0.5 mM NHP induced similar *WRKY38* mRNA accumulation as 1 mM NHP (Supplemental Fig. S2). The abundance of mRNA in the treated seedlings was assessed for each condition via RNA-sequencing (RNA-seq). After general quality control and mapping to the Arabidopsis TAIR10 genome, genes differentially expressed by NHP were determined relative to mock treatment for each time point and genotype using DESeq2 (Love et al., 2014). Genes were considered differentially expressed if they returned an adjusted *P-*value below 0.05 (*P*_adj_ < 0.05).

In wild type seedlings, a total of 2,079 genes were differentially expressed in response to NHP for at least one of the measured time points. To explore the set of early NHP-responsive genes, we first selected for genes showing a robust fold change over mock by applying the cutoff of log_2_(fold change, FC) > 1 for upregulated genes and log_2_(FC) < −1 for downregulated genes. This identified 352 genes that were differentially expressed in response to NHP for at least one time point, with 163 genes up and 189 genes down (Supplemental Table S3). Next, the upregulated genes were hierarchically clustered based on their log_2_(FC) values over the indicated time points and four well-defined gene clusters were identified (Fig. 2A). For each group of genes in the defined clusters, the log_2_(FC) values were then averaged by time point, revealing dynamic patterns of expression. Two of the clusters were upregulated within minutes (designated as ‘early transient’ and ‘early strong’) and two were upregulated by several hours (designated as ‘late weak’ and ‘late strong’) (Fig. 2B, Table 1, Table 2, Supplemental Table S4).

**Figure 2.**
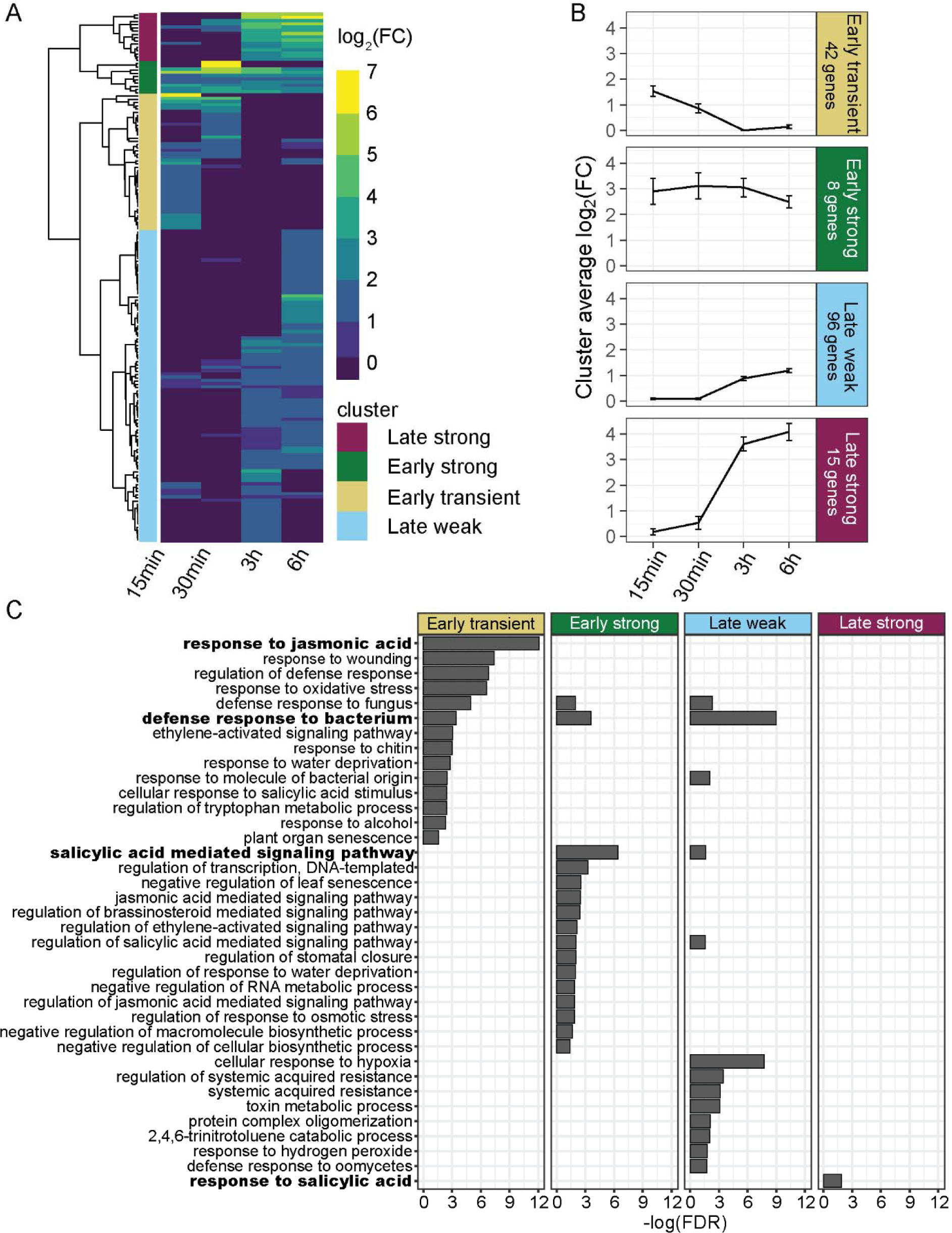
Profile of early NHP-upregulated genes in wild type seedlings. A, Heatmap of genes upregulated (log_2_(FC) > 1, *P*_adj_ < 0.05) in response to NHP treatment in wild type seedlings. One biological replicate consists of 15 pooled seedlings per condition (n = 3). Euclidean distances determined from the log_2_(FC) values were hierarchically clustered across the indicated time points (15 min, 30 min, 3 h and 6 h), resulting in four distinct clusters of gene expression. B, Average log_2_(FC) of all genes within each cluster defined in (A) showing the expression trends across the four time points. Note, the two genes expressed only at 30 min, and no other time, were removed from the ‘early strong’ cluster average. Error bars represent the ± SEM. C, Biological processes of significantly enriched (FDR < 0.05) Gene Ontology (GO) terms for the genes within each cluster are shown. Bars represent the −log(FDR) of each significantly enriched GO term. Bars were omitted if the term was not significantly enriched for the defined cluster. The most significantly enriched GO term for each cluster is indicated in bold.

**Table 1.**
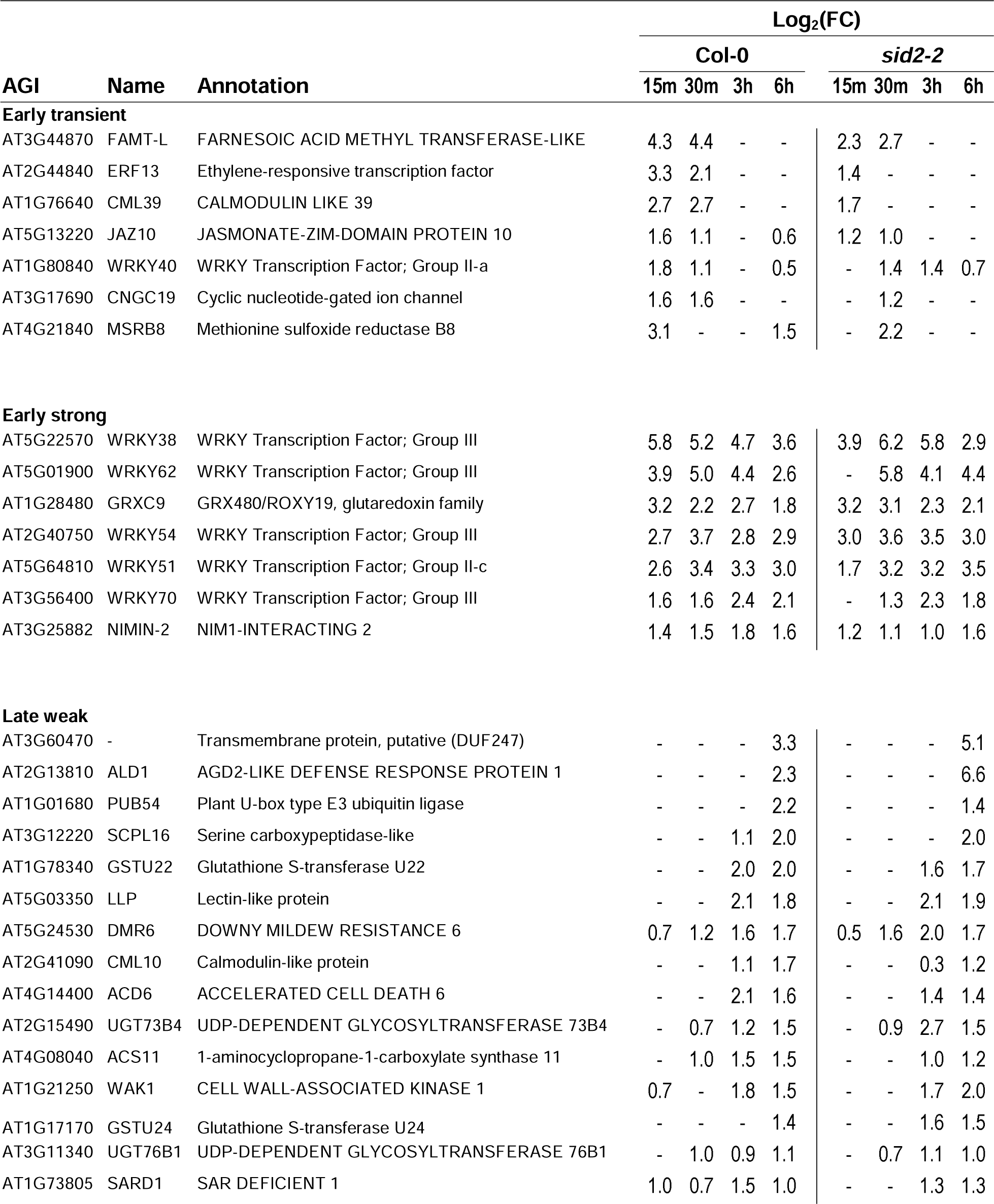
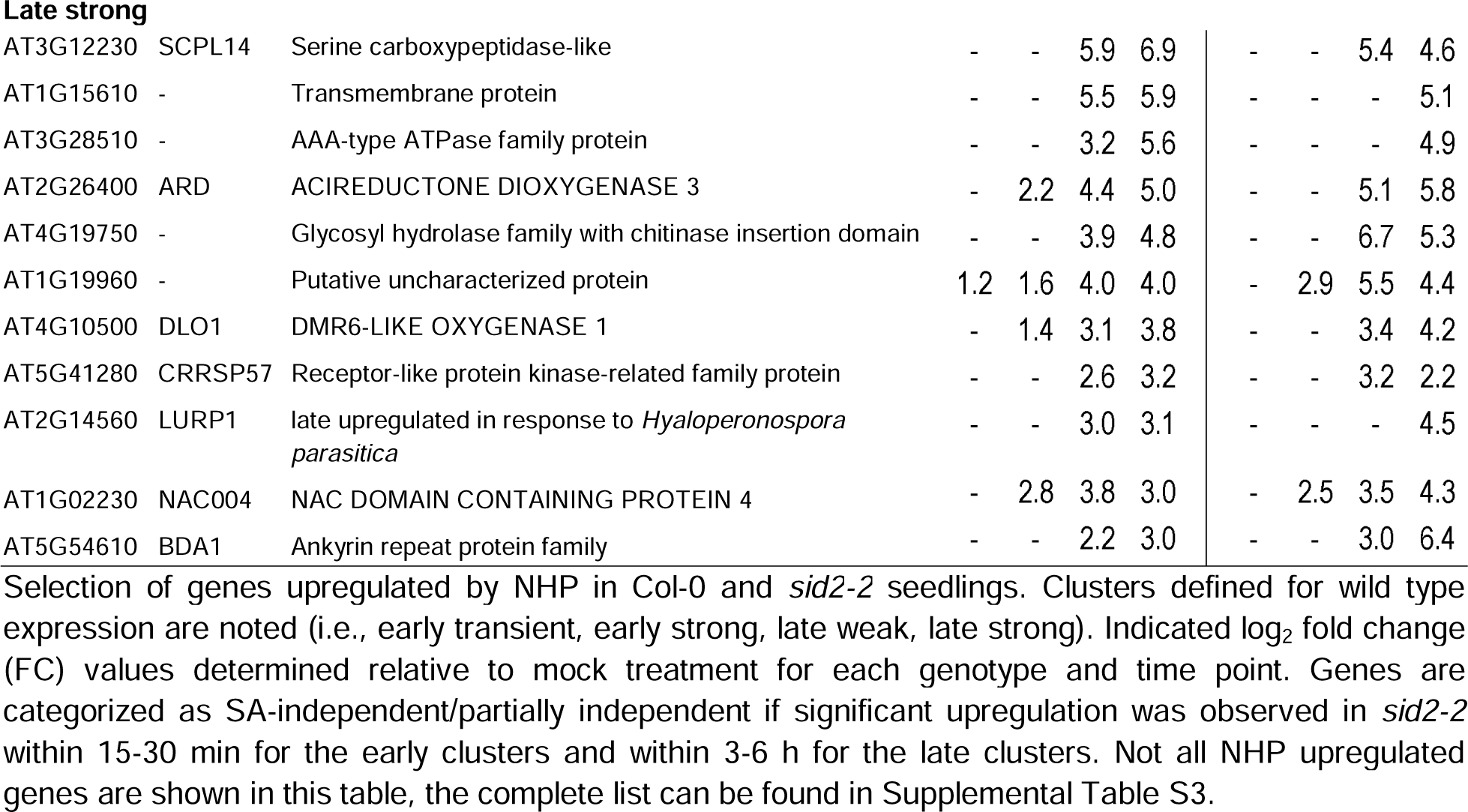
SA-independent/partially independent NHP upregulated genes by cluster.

| AGI | Name | Annotation | Log<sub>2</sub>(FC) |  |  |  |  |  |  |  |
| --- | --- | --- | --- | --- | --- | --- | --- | --- | --- | --- |
|  |  |  | Col-0 |  |  |  | sid2-2 |  |  |  |
|  |  |  | 15m | 30m | 3h | 6h | 15m | 30m | 3h | 6h |
| Early transient |  |  |  |  |  |  |  |  |  |  |
| AT3G44870 | FAMT-L | FARNESOIC ACID METHYL TRANSFERASE-LIKE | 4.3 | 4.4 | - | - | 2.3 | 2.7 | - | - |
| AT2G44840 | ERF13 | Ethylene-responsive transcription factor | 3.3 | 2.1 | - | - | 1.4 | - | - | - |
| AT1G76640 | CML39 | CALMODULIN LIKE 39 | 2.7 | 2.7 | - | - | 1.7 | - | - | - |
| AT5G13220 | JAZ10 | JASMONATE-ZIM-DOMAIN PROTEIN 10 | 1.6 | 1.1 | - | 0.6 | 1.2 | 1.0 | - | - |
| AT1G80840 | WRKY40 | WRKY Transcription Factor; Group II-a | 1.8 | 1.1 | - | 0.5 | - | 1.4 | 1.4 | 0.7 |
| AT3G17690 | CNGC19 | Cyclic nucleotide-gated ion channel | 1.6 | 1.6 | - | - | - | 1.2 | - | - |
| AT4G21840 | MSRB8 | Methionine sulfoxide reductase B8 | 3.1 | - | - | 1.5 | - | 2.2 | - | - |
| Early strong |  |  |  |  |  |  |  |  |  |  |
| AT5G22570 | WRKY38 | WRKY Transcription Factor; Group III | 5.8 | 5.2 | 4.7 | 3.6 | 3.9 | 6.2 | 5.8 | 2.9 |
| AT5G01900 | WRKY62 | WRKY Transcription Factor; Group III | 3.9 | 5.0 | 4.4 | 2.6 | - | 5.8 | 4.1 | 4.4 |
| AT1G28480 | GRXC9 | GRX480/ROXY19, glutaredoxin family | 3.2 | 2.2 | 2.7 | 1.8 | 3.2 | 3.1 | 2.3 | 2.1 |
| AT2G40750 | WRKY54 | WRKY Transcription Factor; Group III | 2.7 | 3.7 | 2.8 | 2.9 | 3.0 | 3.6 | 3.5 | 3.0 |
| AT5G64810 | WRKY51 | WRKY Transcription Factor; Group II-c | 2.6 | 3.4 | 3.3 | 3.0 | 1.7 | 3.2 | 3.2 | 3.5 |
| AT3G56400 | WRKY70 | WRKY Transcription Factor; Group III | 1.6 | 1.6 | 2.4 | 2.1 | - | 1.3 | 2.3 | 1.8 |
| AT3G25882 | NIMIN-2 | NIM1-INTERACTING 2 | 1.4 | 1.5 | 1.8 | 1.6 | 1.2 | 1.1 | 1.0 | 1.6 |
| Late weak |  |  |  |  |  |  |  |  |  |  |
| AT3G60470 | - | Transmembrane protein, putative (DUF247) | - | - | - | 3.3 | - | - | - | 5.1 |
| AT2G13810 | ALD1 | AGD2-LIKE DEFENSE RESPONSE PROTEIN 1 | - | - | - | 2.3 | - | - | - | 6.6 |
| AT1G01680 | PUB54 | Plant U-box type E3 ubiquitin ligase | - | - | - | 2.2 | - | - | - | 1.4 |
| AT3G12220 | SCPL16 | Serine carboxypeptidase-like | - | - | 1.1 | 2.0 | - | - | - | 2.0 |
| AT1G78340 | GSTU22 | Glutathione S-transferase U22 | - | - | 2.0 | 2.0 | - | - | 1.6 | 1.7 |
| AT5G03350 | LLP | Lectin-like protein | - | - | 2.1 | 1.8 | - | - | 2.1 | 1.9 |
| AT5G24530 | DMR6 | DOWNY MILDEW RESISTANCE 6 | 0.7 | 1.2 | 1.6 | 1.7 | 0.5 | 1.6 | 2.0 | 1.7 |
| AT2G41090 | CML10 | Calmodulin-like protein | - | - | 1.1 | 1.7 | - | - | 0.3 | 1.2 |
| AT4G14400 | ACD6 | ACCELERATED CELL DEATH 6 | - | - | 2.1 | 1.6 | - | - | 1.4 | 1.4 |
| AT2G15490 | UGT73B4 | UDP-DEPENDENT GLYCOSYLTRANSFERASE 73B4 | - | 0.7 | 1.2 | 1.5 | - | 0.9 | 2.7 | 1.5 |
| AT4G08040 | ACS11 | 1-aminocyclopropane-1-carboxylate synthase 11 | - | 1.0 | 1.5 | 1.5 | - | - | 1.0 | 1.2 |
| AT1G21250 | WAK1 | CELL WALL-ASSOCIATED KINASE 1 | 0.7 | - | 1.8 | 1.5 | - | - | 1.7 | 2.0 |
| AT1G17170 | GSTU24 | Glutathione S-transferase U24 | - | - | - | 1.4 | - | - | 1.6 | 1.5 |
| AT3G11340 | UGT76B1 | UDP-DEPENDENT GLYCOSYLTRANSFERASE 76B1 | - | 1.0 | 0.9 | 1.1 | - | 0.7 | 1.1 | 1.0 |
| AT1G73805 | SARD1 | SAR DEFICIENT 1 | 1.0 | 0.7 | 1.5 | 1.0 | - | - | 1.3 | 1.3 |
| **Late strong** |  |  |  |  |  |  |  |  |  |  |
| AT3G12230 | SCPL14 | Serine carboxypeptidase-like | - | - | 5.9 | 6.9 | - | - | 5.4 | 4.6 |
| AT1G15610 | - | Transmembrane protein | - | - | 5.5 | 5.9 | - | - | - | 5.1 |
| AT3G28510 | - | AAA-type ATPase family protein | - | - | 3.2 | 5.6 | - | - | - | 4.9 |
| AT2G26400 | ARD | ACIREDUCTONE DIOXYGENASE 3 | - | 2.2 | 4.4 | 5.0 | - | - | 5.1 | 5.8 |
| AT4G19750 | - | Glycosyl hydrolase family with chitinase insertion domain | - | - | 3.9 | 4.8 | - | - | 6.7 | 5.3 |
| AT1G19960 | - | Putative uncharacterized protein | 1.2 | 1.6 | 4.0 | 4.0 | - | 2.9 | 5.5 | 4.4 |
| AT4G10500 | DLO1 | DMR6-LIKE OXYGENASE 1 | - | 1.4 | 3.1 | 3.8 | - | - | 3.4 | 4.2 |
| AT5G41280 | CRRSP57 | Receptor-like protein kinase-related family protein | - | - | 2.6 | 3.2 | - | - | 3.2 | 2.2 |
| AT2G14560 | LURP1 | late upregulated in response to *Hyaloperonospora parasitica* | - | - | 3.0 | 3.1 | - | - | - | 4.5 |
| AT1G02230 | NAC004 | NAC DOMAIN CONTAINING PROTEIN 4 | - | 2.8 | 3.8 | 3.0 | - | 2.5 | 3.5 | 4.3 |
| AT5G54610 | BDA1 | Ankyrin repeat protein family | - | - | 2.2 | 3.0 | - | - | 3.0 | 6.4 |

**Table 2.** SA-dependent NHP upregulated genes by cluster.

| AGI | Name | Annotation | Log<sub>2</sub>(FC) |  |  |  |  |  |  |  |
| --- | --- | --- | --- | --- | --- | --- | --- | --- | --- | --- |
|  |  |  | Col-0 |  |  |  | sid2-2 |  |  |  |
|  |  |  | 15m | 30m | 3h | 6h | 15m | 30m | 3h | 6h |
| Early transient |  |  |  |  |  |  |  |  |  |  |
| AT1G32970 | SBT3.2 | Subtilisin-like protease | 2.8 | 3.2 | - | - | - | - | - | - |
| AT1G21240 | WAK3 | Wall-associated receptor kinase | 2.5 | - | - | 1.4 | - | - | - | 2.0 |
| AT5G44430 | PDF1.2C | PLANT DEFENSIN 1.2C | 2.4 | - | - | - | - | - | - | - |
| AT2G14610 | PR1 | PATHOGENESIS-RELATED GENE 1 | 2.3 | - | - | - | - | - | - | - |
| AT4G34410 | ERF109 | Ethylene-responsive transcription factor | 2.1 | - | - | - | - | - | - | - |
| AT2G32140 | - | Transmembrane receptor | 2.1 | - | - | - | - | - | 2.0 | - |
| AT5G44420 | PDF1.2A | PLANT DEFENSIN 1.2 | 2.0 | - | - | - | - | - | - | - |
| AT3G14260 | - | LURP-one-like protein (DUF567) | 1.8 | - | - | - | - | - | - | - |
| AT2G26020 | PDF1.2B | PLANT DEFENSIN 1.2B | 1.6 | - | - | - | - | - | - | - |
| AT5G64905 | PROPEP3 | ELICITOR PEPTIDE 3 PRECURSOR | 1.5 | - | - | - | - | - | - | - |
| AT1G32640 | MYC2 | JASMONATE INSENSITIVE 1 | 1.4 | - | - | - | - | - | - | - |
| AT2G18660 | EGC2 | PLANT NATRIURETIC PEPTIDE A | 1.4 | - | - | - | - | - | - | - |
| Early strong |  |  |  |  |  |  |  |  |  |  |
| AT2G21900 | WRKY59 | WRKY Transcription Factor; Group II-c | 1.9 | 2.4 | 2.3 | 2.3 | - | - | 2.2 | 2.7 |
| Late weak |  |  |  |  |  |  |  |  |  |  |
| AT2G28850 | CYP710A3 | Cytochrome P450 710A3 | - | - | 2.4 | 2.1 | - | - | - | - |
| AT1G15630 | - | Transmembrane protein | - | - | 1.9 | 1.4 | - | - | - | - |
| AT5G44568 | PROSCOOP4 | SERINE RICH ENDOGENOUS PEPTIDE | - | - | 1.3 | 1.3 | - | - | 0.7 | 1.0 |
| AT1G18830 | SEC31A | Protein transport protein SEC31 homolog A | - | - | - | 2.8 | - | - | - | - |
| AT2G45550 | CYP76C4 | Cytochrome P450 76C4 | - | - | - | 2.6 | - | - | - | - |
| AT1G10155 | ATPP2-A10 | PHLOEM PROTEIN 2-A10 | - | - | - | 2.3 | - | - | - | - |
| AT1G02450 | NIMIN-1 | NIM1-INTERACTING 1 | - | - | - | 2.3 | - | - | - | - |
| AT3G45860 | CRK4 | Cysteine-rich receptor-like protein kinase | - | - | - | 2.1 | - | - | - | - |
| AT5G51500 | PME60 | Pectin methylesterase inhibitor | - | - | - | 1.7 | - | - | - | - |
| AT2G29350 | SAG13 | SENESCENCE-ASSOCIATED GENE 13 | - | - | - | 1.3 | - | - | - | - |
| Late strong |  |  |  |  |  |  |  |  |  |  |
| AT3G22231 | PCC1 | PATHOGEN AND CIRCADIAN CONTROLLED 1 | - | - | 3.8 | 3.9 | - | - | - | - |

The ‘early transient’ cluster includes 42 genes and is defined by elevated transcript abundance 15-30 min after NHP treatment that did not persist by 3 h. Gene ontology (GO) analysis for this cluster showed strong enrichment for biological processes associated with responses to jasmonic acid (JA), wounding, and oxidative stress. Some of these genes are associated with the regulation of defense as well as defense responses to fungi and bacteria (Fig. 2C). Among the most highly expressed genes in this cluster are JA-responsive genes, which play key roles in the regulation of wounding responses, including *CML39* which encodes a calmodulin-like protein, *JAZ10* which encodes a jasmonate-zim-domain protein, and *MYC2* which encodes a jasmonate-inducible transcription factor (Table 1, Table 2). Taken together, the early yet unsustained induction of these genes within 30 min suggests a burst of JA- and wound-responsive gene expression that acts as a primary wave of NHP signaling.

The ‘early strong’ cluster includes 10 genes and is defined by a high average log_2_(FC) 15 min after NHP treatment that was maintained for all subsequent time points. Of the 10 genes, two (AT1G06475 and AT5G40980) were strongly upregulated by NHP at 30 min and no other time points. These two genes were removed from the average log_2_(FC) analysis and subsequent analyses because they did not behave like the other genes in this cluster (Fig. 2B). The early strong cluster is notably enriched in biological processes involved in the regulation of transcription, with over half the genes belonging to the WRKY TF family, including four group III *WRKY* genes (*WRKY38*, *WRKY54*, *WRKY62* and *WRKY70*) and two group II-c genes (*WRKY51* and *WRKY59*) (Eulgem et al., 2000; Kalde et al., 2003); (Table 1, Table 2, Supplemental Table S4). Additional GO terms enriched in the early strong cluster includes regulation of defense and regulation of hormonal signaling pathways such as SA, JA, brassinosteroid, and ethylene (Fig. 2C). The early and sustained induction of these genes, particularly the *WRKYs*, suggests they are primary transcriptional target genes of NHP signaling.

The ‘late weak’ cluster includes the largest number of genes (96) that averaged a log_2_(FC) around 1 (Fig. 2B). GO term analysis of the cluster revealed significant enrichment of biological processes involved in response to bacteria and oomycetes, as well as SA-mediated signaling and regulation of SAR (Fig. 2C). This indicates NHP activation of SAR and pathogen defense genes can be detected as early as 3-6 h in treated tissues. Notably, genes involved in the regulation and biosynthesis of NHP were observed in this wave of transcription, including *SARD1* (*SAR DEFICIENT 1*), *NIMIN-1* (*NIM-INTERACTING 1*), *ALD1,* and *UGT76B1* (Table 1, Table 2, Supplemental Table S4) (Song et al., 2004; Zhang et al., 2010a; Holmes et al., 2021).

The ‘late strong’ cluster consists of 15 genes with log_2_(FC) values averaging between 3.5 - 4 (Fig. 2B). Response to SA was the only significantly enriched GO term for this cluster (Fig. 2C). Despite having a role in the response to SA, the majority of genes classified as late strong were upregulated by NHP in both wild type and *sid2-2* (Fig. 3A, Table 1), indicating that accumulation of these transcripts was at least partially SA-independent. Of the 14 SA-independent genes identified in this cluster, only five have been found to be upregulated 24 h after NHP treatment in an SA-independent manner (Yildiz et al., 2021). Thus, our analysis has uncovered additional NHP-responsive genes that operate independently of SA as early as 6 h post NHP signaling.

**Figure 3.**
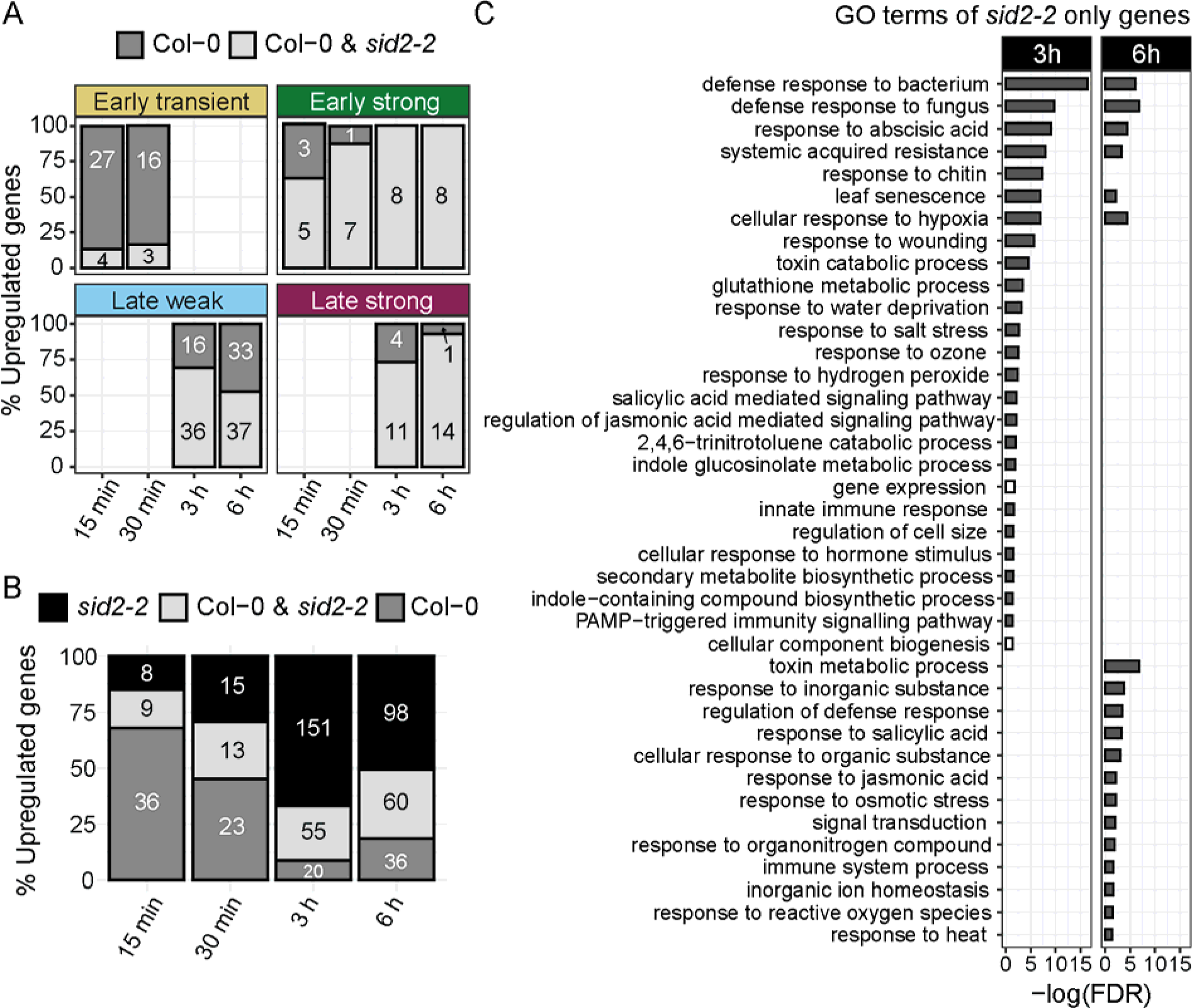
Comparison of NHP upregulated genes in wild type and *sid2-2* seedlings. A, Percent of NHP upregulated genes expressed only in wild type (Col-0, SA-dependent) and in both wild type and the *sid2-2* mutant (SA-independent) for the wild type gene expression clusters defined in Figure 2. Total number of genes in each condition is indicated. B, Total number of NHP upregulated genes (log_2_(FC) > 1, *P*_adj_ < 0.05) per time point unique to wild type (Col-0), shared between wild type and *sid2-2*, and unique to *sid2-2*. The number of upregulated genes in each grouping is indicated. C, Biological processes of significantly enriched (gray bar) or depleted (white bar) (FDR < 0.05) GO terms for the set of genes upregulated 3 and 6 h after NHP treatment that were upregulated only in the *sid2-2* mutant and not wild type. Bars represent the negative log(FDR) of each significantly enriched GO term, bars were omitted if the term was not significantly enriched or depleted for the defined time point.

We also performed a similar clustering analysis on the 189 NHP-downregulated transcripts and identified three gene clusters (Supplemental Fig. S3A, B). In contrast to the upregulated gene clusters, the downregulated gene clusters had less distinct patterns of expression and averaged log_2_(FC) values around −1 to −1.5. Two clusters were downregulated within 15-30 min and persisted 3-6 h hours after NHP treatment. One of these showed weaker downregulation and was classified as an ‘early weak’ cluster. The other showed stronger downregulation and was classified as an ‘early strong’ cluster. The third did not show decreased transcript abundance until 3-6 h after treatment and thus was labeled as a ‘late’ cluster (Supplemental Fig. S3B). All three clusters were enriched for biological processes involving growth and development such as root morphogenesis, plant epidermis development, plant-type cell wall organization, and root hair cell differentiation (Supplemental Fig. S3C). Given the NHP precursor Pip is known to inhibit root growth of seedlings (Wang et al., 2018), it is likely that NHP also inhibits root growth. These clusters shed new light on the genes that may be associated with altered plant growth and development in response to elevated levels of NHP.

### SA biosynthesis is dispensable for the early induction of a majority of NHP upregulated genes

Interrogation into the late strong cluster suggested NHP may be sufficient to activate a subset of SA-responsive genes in the absence of NHP-induced SA biosynthesis (Fig. 2C, Fig. 3A, ‘late strong’). This was an intriguing possibility given NHP likely accumulates in distal tissues prior to SA accumulation following a local tissue infection (Bernsdorff et al., 2016; Hartmann et al., 2018; Hartmann and Zeier, 2019). We therefore hypothesized that NHP would need to drive transcriptional reprogramming as SA levels rise from low to high. To investigate this, we assessed the impact of SA biosynthesis on NHP-induced gene expression from 15 min to 6 h. For each time point of the wild type gene clusters described in Fig. 2, the percentage of genes upregulated in both wild type and the *sid2-2* mutant was determined. This group of genes was defined as ‘SA-independent’. Genes upregulated only in wild type and not in *sid2-2* at the defined time point were conversely defined as ‘SA-dependent’.

Notably, the majority of genes upregulated by NHP at 15 min in the early transient cluster were SA-dependent. For example, 87% of the genes (27 out of 31) were upregulated at 15 min in wild type but not in *sid2-2* seedlings (Fig. 3A) and similar trends were seen at 30 min. By contrast, 100% of the genes (8 out of 8) in the early strong cluster upregulated 3-6 h after NHP treatment were SA-independent (Fig. 3A). 69% of the genes (36 out of 52) in the late weak cluster were SA-independent at 3h, which dropped to 53% (37 out of 70 genes) at 6 h (Fig. 3A).

A similar pattern of gene expression was observed when we analyzed the total number of NHP upregulated genes reaching a log_2_(FC) > 1 without gene clustering (Fig. 3B). The majority of genes induced by NHP between 15-30 min were SA-dependent (i.e., 36 genes up in wild type and 9 up in both wild type and *sid2-2* at 15 min) and the majority of genes induced by NHP between 3-6 h were SA-independent (i.e., 20 genes up in wild type and 55 up in both wild type and *sid2-2* at 3 h) (Fig. 3B). Taken together, this analysis highlights an important temporal role of SA on NHP induced transcription. Our data suggest that new SA biosynthesis is required for the induction of NHP-responsive genes within minutes of exposure to NHP, whereas SA biosynthesis is dispensable for NHP-induced expression several hours later.

We also analyzed the downregulated genes using the same approach and found a slight trend of SA-dependence for these genes following NHP treatment. For example, the early weak and late clusters defined in Supplemental Fig. S3 showed that a majority of genes are downregulated in wild type but not the *sid2-2* mutant at each time point (Supplemental Fig. S4A). By contrast, the genes of the early strong cluster were less dependent on SA biosynthesis, such that by the 6 h time point only 15% of genes (8 out of 44) were downregulated in wild type and not in *sid2-2* seedlings (Supplemental Fig. S4A). Independent of clustering, greater than 60% of the downregulated genes were found to be SA-dependent at 15 min and 3 h and approximately 40% were SA-dependent at 30 min and 6 h (Supplemental Fig. S4B). Taken together, these findings suggest a stronger dependence on SA biosynthesis for NHP-dependent downregulation of gene expression over the time courses studied.

### SA biosynthesis antagonizes NHP-elicited gene expression at 3 to 6 h

Looking at the genes upregulated only in the *sid2-2* mutant and not wild type, we discovered an unexpectedly large number of transcripts that increased in abundance upon NHP treatment at 3 and 6 h post treatment (Fig. 3B). This includes 151 genes at 3 h and 98 genes at 6 h that were significantly upregulated by NHP in the *sid2-2* mutant but not in wild type (Fig. 3B). GO-term enrichment analysis of these genes showed significant enrichment for a number of biotic and abiotic stress responses, such as defense response to bacteria and fungi, SAR, response to abscisic acid, and response to hypoxia (Fig. 3C). These data suggest that SA may play an important role in modulating the transcription of a suite of genes, through an unknown mechanism, once NHP-mediated transcriptional reprogramming has been initiated.

### TGA and WRKY transcription factor *cis*-regulatory elements are enriched in the promoters of early NHP upregulated genes

To identify TFs that might directly control the expression of early NHP-responsive genes, we analyzed the promoter regions of all genes upregulated after NHP treatment in both wild type and *sid2-2* mutant seedlings to identify putative TF bindings sites (i.e., *cis-*regulatory elements, CREs). A less stringent cutoff of log_2_(FC) > 0 and *P*_adj_ < 0.05 was used for the upregulated genes to capture putative transcriptional regulators of all genes significantly elevated in transcript abundance, regardless of the magnitude of expression change. Following a modified workflow detailed by Bjornson et al., 2021, the NHP upregulated genes were then separated by the time points in which they were differentially expressed. Enrichment of transcription factor CREs was then determined for each time point using the DAPseq database (O’Malley et al., 2016) (Fig. 4A). Supplemental Fig. 5 shows all the CREs found to be significantly enriched in the promoters of NHP-upregulated genes in both wild type and *sid2-2* seedings between 15 min and 6 h following NHP treatment.

**Figure 4.**
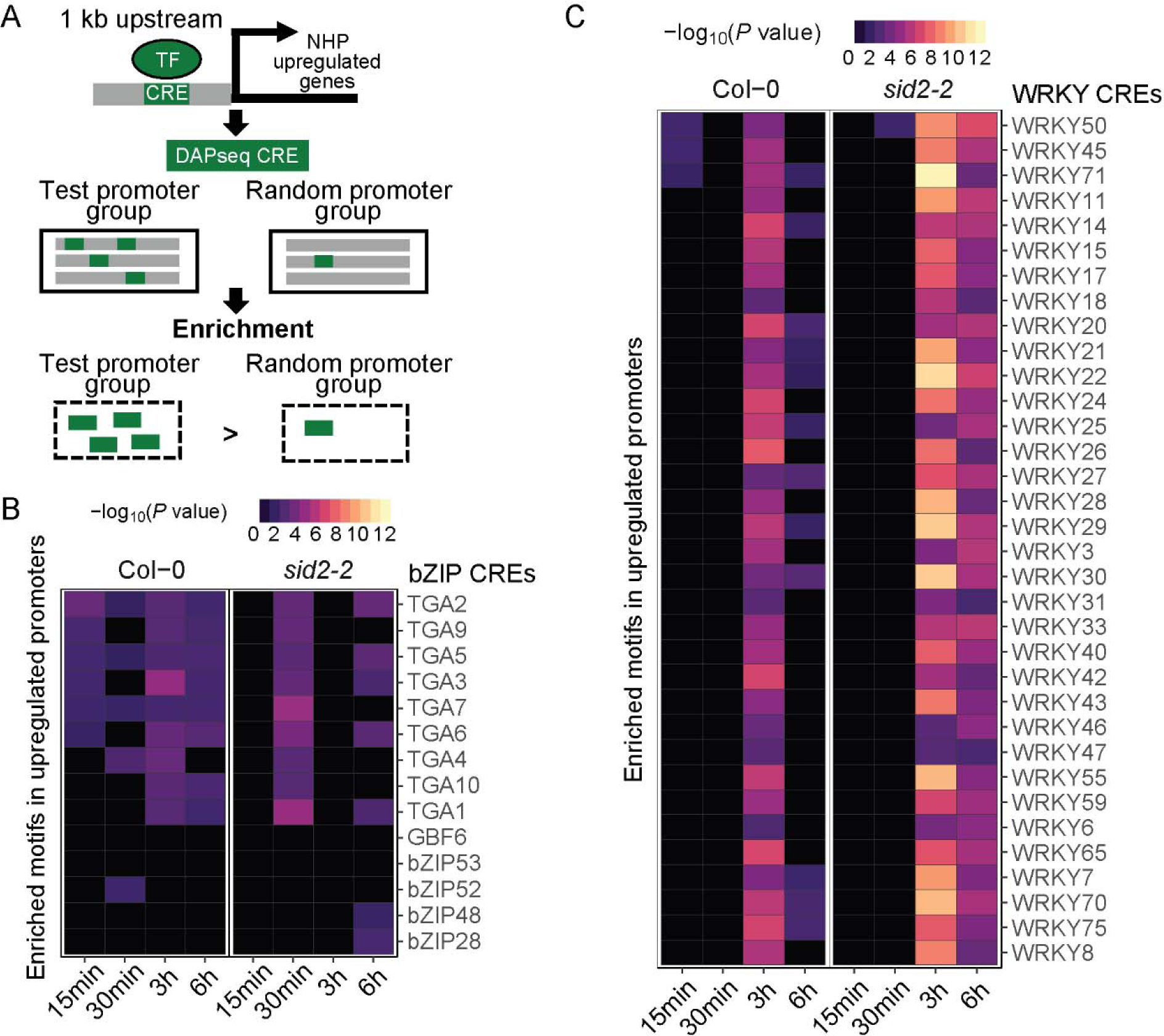
Presence of TGA/bZIP and WRKY TF *cis*-regulatory elements (CREs) in the promoters of NHP-upregulated genes in wild type (Col-0) and *sid2-2* seedlings. A, Schematic of the TF binding motif (i.e., CRE) enrichment analysis performed. All genes with increased transcript abundance upon NHP treatment (log_2_(FC) > 0 and *P*_adj_ < 0.05) were grouped by time point for the indicated genotype, these groups are represented as “test promoter group” in the diagram. CREs of known Arabidopsis TFs from the DAPseq database (O’Malley et al., 2016) were identified in the promoters (1 kb upstream of the transcriptional start site) of each group of promoters. Enrichment of identified CREs was determined relative to CREs found in promoters pulled from a random sampling of genes detected in this RNA-seq experiment. Schematic design is modified from Mariani et al., 2017. Enrichment analysis of TGA/bZIP (B) and WRKY (C) CREs. The name of the TF known to bind the enriched CRE is listed on the right and the TF family name is indicated above (i.e., bZIP or WRKY). The −log_2_(*P* value) of the enrichment analysis is indicated by the scale bar with black indicating no significant enrichment of the CRE and purple to yellow denoting an enriched CRE.

We found that CREs for bZIP family members, specifically TGA TFs, were enriched in the promoters of genes upregulated by NHP as early as 15 min. For some TGAs (e.g., TGA2, TGA5, and TGA7), the enrichment of their TF binding sites was observed for all time points (Fig. 4B). Enrichment of TGA binding sites in the promoters of early NHP upregulated genes is consistent with previous work demonstrating TGA2/5/6 and TGA1/4 are required for the induction of NHP responsive genes at 24-48 h post NHP treatment (Nair et al., 2021; Yildiz et al., 2023). TGA TFs are known to interact with the transcriptional coactivator NPR1 to drive SAR- and SA-induced transcriptional changes (Zhang et al., 1999). The enrichment of TGA binding sites in the promoters of NHP upregulated genes as early as 15 min implicates TGAs and NPR1 among the primary transcriptional regulators of NHP signaling. Furthermore, enrichment of TGA CREs was not observed in *sid2-2* seedlings until 30 min and was also not significantly enriched at 3 h (Fig. 4B). These findings suggest that SA biosynthesis and likely elevated SA levels are required for rapid and stable induction of some TGA-regulated genes induced by NHP signaling.

Moreover, we discovered that a large number of WRKY CREs were highly enriched in the promoter regions of NHP upregulated genes at 3-6 h in both wild type and *sid2-2* seedlings (Fig. 4C). This correlates with the majority of genes identified in the NHP upregulated early strong cluster belonging to the WRKY TF family (Fig. 2, Table 1). This finding suggests a model where NHP induces the expression of *WRKY* genes as a primary transcriptional response, followed by WRKY-regulated gene expression changes that act as a secondary wave of transcriptional control.

It is striking that the confidence in WRKY CRE enrichment was greater (i.e., lower *P* value) at 3 and 6 h in the promoters of genes upregulated in the *sid2-2* mutant compared to those of wild type (Fig. 4C). These data suggest that a substantial number of NHP upregulated genes are directly regulated by WRKY TFs, and such regulation is influenced by SA levels. A similar pattern emerged for members of the NAC TF family, where NAC CREs were enriched in the promoters of NHP upregulated genes at 3 h in *sid2-2* but not in the promoters of genes upregulated in wild type (Supplemental Fig. S5). Both WRKY and NAC TFs are involved in the regulation of a number of biotic and abiotic stress responses (Eulgem et al., 2000; Mohanta et al., 2020). Notably, the set of genes upregulated by NHP at 3-6 h only in the *sid2-2* mutant but not in wild type were enriched in biological processes associated with biotic and abiotic stress responses (Fig. 3B, C). Taken together, these data suggest SA may antagonize the transcription of a subset of NHP responsive genes by regulating the activities of WRKY and NAC TFs.

The same CRE enrichment analysis was also carried out on all NHP downregulated genes, regardless of the magnitude of fold change (log_2_(FC) < 0 and *P*_adj_ < 0.05). We found multiple WRKY CREs enriched in the promoter regions of NHP downregulated genes at 15 min, 3 h, and 6 h, in wild type seedlings (Supplemental Fig. S6), further supporting a role for WRKY TFs in relaying NHP-induced transcriptional changes.

Finally, we found the CREs of Homeobox gene family members to be abundant in the promoters of NHP downregulated genes spanning from 15 min through 6 h, with the greatest confidence of enrichment found in the promoters of genes downregulated by NHP at 3-6 h in an SA-dependent manner (Supplemental Fig. S6). Among these was ATHB5, a homeodomain leucine zipper (HDZip) protein linked to the abscisic acid (ABA)-driven repression of germination and root growth in seedlings (Johannesson et al., 2003). Given that NHP precursor Pip inhibits seedling root growth, we now hypothesize that NHP may drive the repression of root growth and development genes (Supplemental Fig. 3C) through the action of HDZip Homeobox proteins like ATHB5.

### Early NHP-responsive gene *WRKY70* is required for NHP-elicited SAR

To further dissect the NHP response pathway, we carried out a reverse genetic screen to identify Arabidopsis mutants with compromised NHP-elicited SAR. Given WRKY CREs were the predominant regulatory elements identified in genes responsive to NHP 3-6 h following treatment (Fig. 4C, Supplemental Fig. S6), we selected the six *WRKY*s (*WRKY38, WRKY51, WRKY54, WRKY59, WRKY62, WRKY70)* from the early strong NHP upregulated cluster (Figure 2B, Table 1, Table 2) for mutant analysis.

To measure NHP-induced SAR, we obtained homozygous Arabidopsis mutants and quantified bacterial growth in their leaves after treatment with exogenous NHP (Chen et al., 2018). Specifically, three lower leaves of 4.5-week-old wild type and mutant plants were infiltrated with water (mock) or 0.5 mM NHP. One day later, one distal upper leaf was inoculated with a 1 x 10^5^ CFU/mL suspension of the virulent bacterium *Pseudomonas syringae* pathovar *maculicola* strain ES4326 (*Psm*). Bacterial growth of *Psm* was quantified three days post infection (dpi) to determine if the mutants were altered in their resistance to pathogen infection. Mutants were considered to exhibit NHP-elicited SAR if bacterial titer was significantly lower in NHP treated plants compared to mock treated plants. By contrast, mutants were considered insensitive or partially insensitive to NHP if the bacterial titer of NHP treated mutants was higher than NHP treated wild type plants.

Of the six *wrky* mutants screened, five (*wrky38*, *wrky62*, *wrky54*, *wrky51*, and *wrky59*) exhibited normal NHP-elicited SAR (Fig. 5B-E). Only one, *wrky70-*1 (T-DNA insertional line SALK_025198) (Li et al., 2006), displayed compromised NHP-elicited SAR (Fig. 5A). That is, NHP treatment did not decrease bacterial growth in distal leaves of *wrky70-1* plants compared to mock treatment. We also observed that *wrky70-1* mutants treated with mock contained less bacteria than wild type plants treated with mock, demonstrating that basal resistance is enhanced in the *wrk70-1* mutant (Fig. 5A). Notably, distal leaves of *wrky70-1* plants treated with NHP contained higher levels of bacteria than distal leaves of wild type plants treated with NHP (Fig. 5A). The inability of *wrky70-1* plants to further restrict bacterial growth following NHP treatment suggests that the mutant may be insensitive to NHP or defective in NHP signaling. Taken together, these data indicate that *WRKY70* is required for full NHP-elicited systemic resistance.

**Figure 5.**
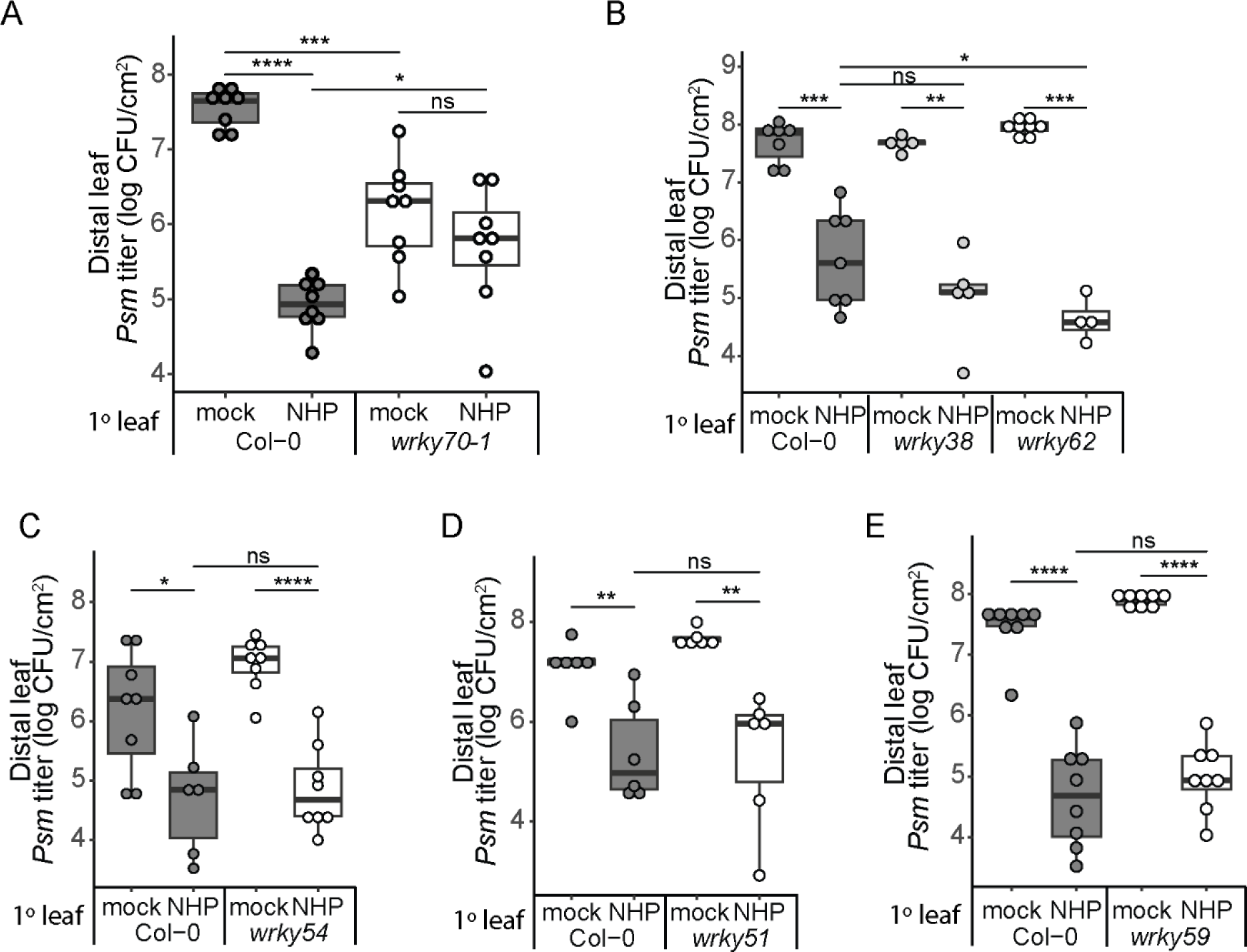
*WRKY70* is involved in NHP-elicited SAR. Bacterial growth in the distal leaves of mock and NHP treated wild type (Col-0) and *wrky* mutant plants. *WRKY* genes for mutant analysis were selected from the early stable cluster defined in Figure 2 and include *WRKY70* (A), *WRKY38* and *WRKY62* (B), *WRKY54* (C), *WRKY51* (D), and *WRKY59* (E). Three lower (1°) leaves were infiltrated with water (mock), 0.5 mM NHP, or 1 mM NHP (only for *wrky59*) and 1 d later one upper, distal leaf was inoculated with a 1 x 10^5^ CFU/mL suspension of *Psm*, followed by quantification of bacterial titer 3 dpi (n = 4-8). Asterisks indicate significant differences in bacterial titer (two-tailed t-test; \**P* < 0.05, \*\**P* < 0.01, \*\*\**P* < 0.001, \*\*\*\**P* < 0.0001, ns = not significant).

We next questioned if systemic resistance to *Psm* was similarly compromised in *wrky70-1* plants primed with the avirulent pathogen *P*. *syringae* pathovar *tomato* (*Pst*) strain DC3000 carrying *avrRpt2* (*Pst avrRpt2*). Localized infection with *Pst avrRpt2* is known to elicit a strong immune response that leads to SAR and protection against *Psm* in distal leaves (Kohler et al., 2002). We found that the *Psm* titer in distal leaves reached similar levels for *wrk70-1* plants treated with mock or *Pst avrRpt2* and this level was comparable to *Psm* growth in wild type leaves of plants treated with *Pst avrRpt2* (Supplemental Fig. S7A). These data indicate that *wrky70-1* mutants exhibit a basal-level of defense priming and *Pst avrRpt2* primary infection does not further enhance this resistance.

We also examined pathogen growth in local *wrky70-1* leaves. We found that the titer of *Psm* in infected *wrky70-1* leaves was lower than in *Psm* infected wild type leaves (Supplemental Fig. S7B). This phenotype corroborates previous work demonstrating loss of *WRKY70* expression enhances resistance to *Psm* infection (Zhou et al., 2018).

### Expression of *SARD1* and *PR* genes is elevated in *wrky70-1* independent of NHP

Previous studies in Arabidopsis have demonstrated a role for WRKY70 in the transcriptional regulation of several genes required for SAR signaling and disease resistance, including *SARD1* and the *PATHOGENESIS-RELATED* genes *PR1*, *PR2*, and *PR5* (Li et al., 2004; Zhou et al., 2018; Liu et al., 2021). These genes were also shown to be upregulated in wild type Arabidopsis plants 24 h after treatment with NHP (Yildiz et al., 2021). To determine if WRKY70 is required for NHP induced expression of *SARD1*, *PR1*, *PR2*, and *PR5* at 24 h, we quantified transcript abundance in wild type and *wrky70-1* plants treated with water or 1 mM NHP. As expected, qRT-PCR showed increased abundance of *SARD1*, *PR1*, *PR2*, and *PR5* mRNA in wild type plants treated with NHP compared to water (Supplemental Fig. S7C). However, NHP treatment did not further increase transcript levels in *wrky70-1* plants (Supplemental Fig. S7C). Notably, all four genes showed elevated transcript abundance in water treated *wrky70-1* mutants compared to water treated wild type. The elevated expression of *SARD1* and *PR1* in water treated *wrky70-1* plants is consistent with previous studies demonstrating WRKY70 negatively regulates these two genes in the absence of a pathogen (Zhou et al., 2018; Liu et al., 2021).

### WRKY70 is required for NHP induction of genes in the late weak and late strong clusters

Given *wrky70-1* mutants exhibited enhanced basal resistance but were still unable to achieve the same level of resistance as NHP treated wild type plants (Figure 5A), we hypothesized that a subset of early NHP-responsive genes containing putative WRKY70 CREs in their promoters may be dependent on WRKY70 function for full, wild type gene expression levels following NHP treatment. To investigate this, we selected three genes from the late weak and late strong clusters (Fig. 2, Table 1), specifically *BDA1*, *PROSCOOP4*, and *CML10,* and examined their transcript accumulation in response to 1 mM NHP over a time course of 0 to 12 h. The promoters of these genes contain one or more WRKY70 CREs (i.e., W-box (TTGACY), WT-box (YGACTTTT), and WRKY70 DAPseq motif; (Rushton et al., 2010; Machens et al., 2014; O’Malley et al., 2016)) within 2000 base pairs upstream of their respective putative transcriptional start site (Fig. 6A).

**Figure 6.**
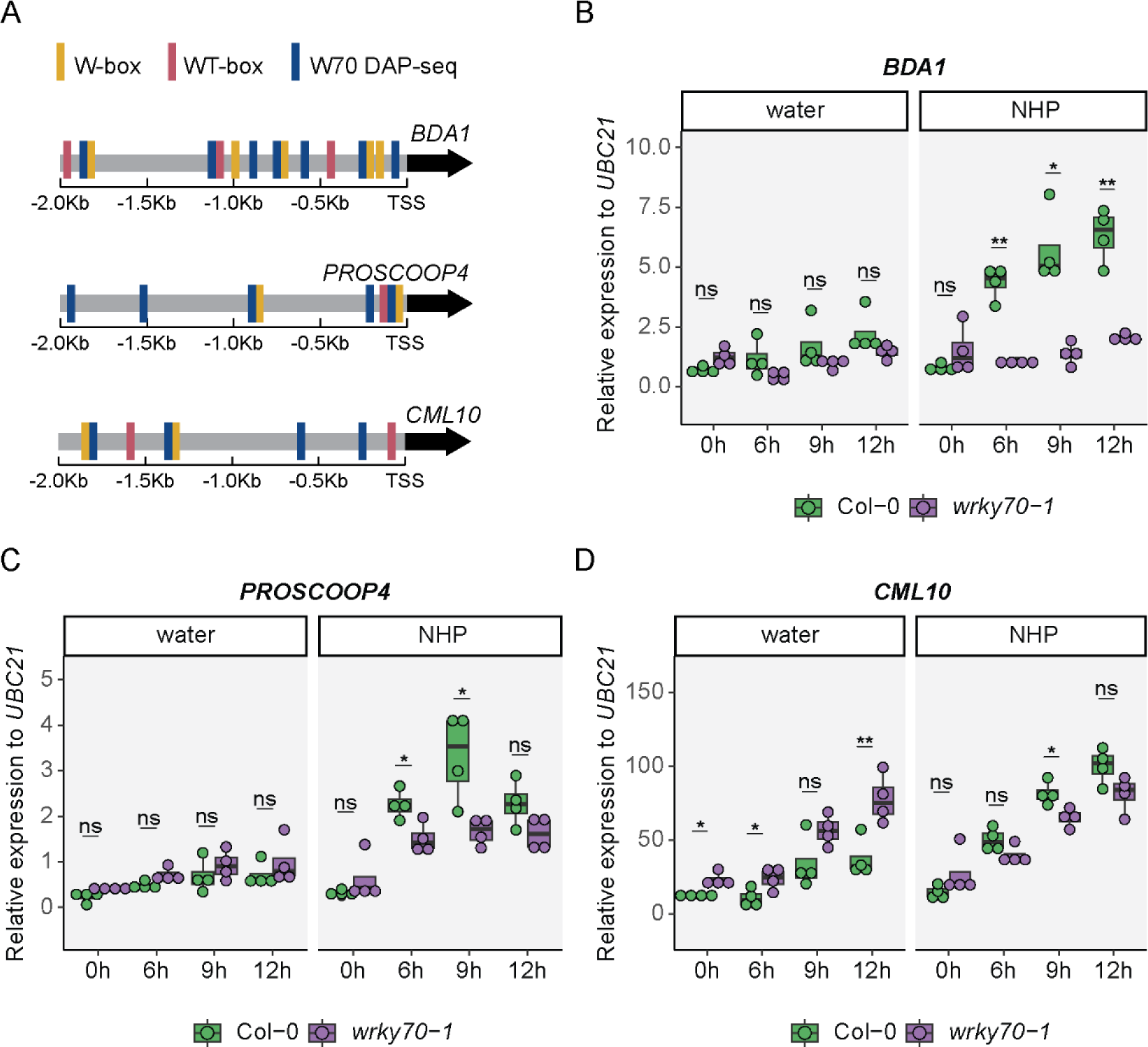
Loss of WRKY70 function impacts upregulation of early NHP-responsive genes. A, Diagram of putative WRKY70 CREs, including W-box, WT-box, and WRKY70 (W70) DAPseq motifs in the promoters (−2 Kb upstream of the transcriptional start site; TSS) of NHP-responsive genes *BDA1*, *PROSCOOP4*, and *CML10*. Note W70 DAPseq sites adjacent to a W- or WT-box overlap. Expression of *BDA1* (B), *PROSCOOP4* (C), and *CML10* (D) in *wrky70-1* mutant plants. Three leaves of 4.5-week-old wild type (Col-0) and *wrky70*-1 mutant plants were infiltrated with water or 1 mM NHP. Samples collected at 0 h were untreated before collection. Transcript abundance was determined relative to *UBC21* for each condition (2^−ΔCt^). Asterisks indicate a significant difference between wild type and *wrky70-1* at each time point (two-tailed t-test; \**P* < 0.05, \*\**P* < 0.01, ns = not significant).

We found that *BDA1* (*bian da*; “becoming big” in Chinese), a gene encoding an ankyrin-repeat transmembrane protein (Yang et al., 2012), exhibited the strongest dependence on WRKY70 for NHP-induced expression. *BDA1* transcripts were significantly increased at 6, 9, and 12 h post NHP treatment in wild type leaves but not *wrk70-1* leaves (Fig. 6B). Transcript levels for *PROSCOOP4*, a gene encoding the precursor of secreted peptide SCOOP4/STMP10 (Gully et al., 2019; Hou et al., 2021), were also significantly higher at 6 and 9 h post NHP treatment in wild type leaves compared to that observed for *wrk70-1* plants (Fig. 6C). Unlike *BDA1*, we found that *PROSCOOP4* transcripts were higher in NHP-treated *wrk70-1* leaves at 6 and 9h compared to the 0h timepoint (Fig. 6C). These data show that *wrk70-1* plants responded to NHP treatment, suggesting that they can sense NHP but are impaired in NHP signaling. Similar trends were observed for *CML10* (*CALMODULIN-LIKE 10*/*CaBP22*; (Luan et al., 2002)) at the 9 h timepoint (Fig. 6D). We confirmed these results by analyzing a second *wrky70* allele, *wrky70-2* (Supplementary Fig. S8).

Taken together, our findings reveal that *WRKY70* is required for the proper expression of NHP responsive genes from the late weak and late strong clusters, highlighting a role for WRKY70 in the transcriptional response to NHP several hours following treatment.

### NHP pretreatment enhances flg22-elicited ROS production and WRKY70 contributes to the full response

We next questioned if WRKY70 regulates a specific branch of NHP defense signaling. One such branch involves the response to common molecular features of microbes (e.g., bacterial flagellin or fungal chitin) known as microbe-associated molecular patterns (MAMPs) and the response to plant-derived damage-associated molecular patterns (DAMPs), a process collectively referred to as pattern-triggered immunity (PTI). The WRKY70-dependent gene *BDA1* is known to mediate signaling in response to MAMP detection and interacts with receptor-like protein SNC2 (Suppressor of NPR1, Constitutive2) to relay MAMP-triggered defense responses (Yang et al., 2012). Production of reactive oxygen species (ROS) is a MAMP-triggered defense response which can act directly or indirectly as an antimicrobial agent and serves as a secondary signal to activate further defense responses (Boller and Felix, 2009; Couto and Zipfel, 2016). Similarly, DAMPs, such as secreted peptides from the PROSCOOP family, are known to elicit an increase in ROS production (Gully et al., 2019; Hou et al., 2021; Rhodes et al., 2021). Incidentally, *CML10* encodes a calmodulin-like protein known to interact with phosphomannomutase (PMM) in order to modulate ascorbic acid synthesis and cellular homeostasis of ROS (Cho et al., 2016).

We thus investigated how NHP alters MAMP-triggered ROS production using bacterial flagellin as the elicitor in the presence and absence of NHP pretreatment and asked if WRKY70 is required for the response. Leaf discs of wild type and *wrky70-1* plants were floated on solutions of water, NHP, and SA. SA was included as a positive control, as it is known to enhance the MAMP-triggered ROS burst (Yi et al., 2014). After 24 h, leaf discs were treated with a 100 nM solution of flagellin peptide (flg22) and then ROS was measured using a luminol-based assay. We found ROS levels from mock treated *wrky70-1* plants trended lower than water treated wild type, suggesting WRKY70 is required for maximal ROS accumulation (Fig. 7, upper panel). NHP treatment increased ROS generation in both wild type and *wrky70-1* tissue exposed to flg22 when compared to water treated tissues. The average ROS production in NHP treated *wrky70-1* leaf discs was lower than NHP treated wild type leaf discs (Fig. 7, lower panel). Similar results were found when analyzing *wrky70-2* plants (Supplementary Fig. S9). Collectively, these data indicate that NHP is able to enhance the flg22-elicited ROS burst, and this enhancement requires WRKY70 for full ROS generation.

**Figure 7.**
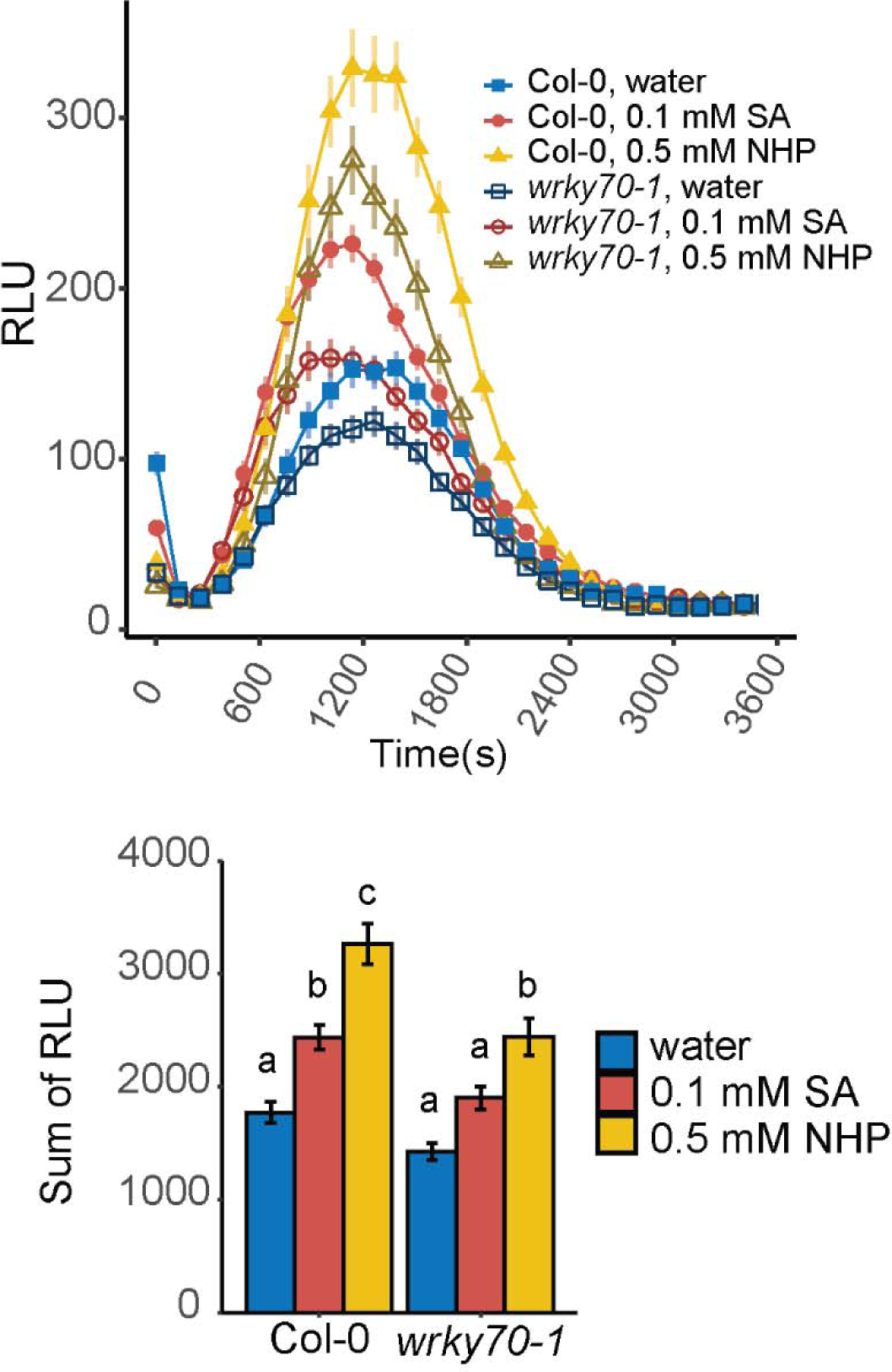
NHP enhances ROS production and requires WRKY70 for full ROS levels. Flg22-elicited ROS quantification in *wrky70-1* plants. Four leaf discs from 4.5 to 5-week-old wild type and *wrky70-1* plants were pretreated by floating leaf discs on water, 0.1 mM SA, or 0.5 mM NHP for 24 h before treating with 100 nM flg22 in horseradish peroxidase and Luminol. Top, traces of the average relative luminescence units (RLU) over the indicated run time. Each point represents the average of 6 plants (n = 24) from two experiments. Bottom, total quantification of RLUs from top plot. For each leaf disc, RLUs from each time point were summed by condition over the run time and averaged with error bars representing the standard error of the mean. Statistical analysis was performed using a one-way ANOVA and post hoc Sidak test, different letters indicated statistical differences between means with *P* < 0.05.

## Discussion

NHP has emerged as an essential signaling molecule required for the establishment and amplification of SAR in Arabidopsis and crop plants (Chen et al., 2018; Holmes et al., 2019), though the early signaling events mediated by NHP had yet to be described. Our study provides the first insights into the primary transcriptional responses to NHP, as well as the role that NHP-induced SA biosynthesis plays during this response. We also discovered that TGA and WRKY CREs are enriched in the promoters of early NHP induced genes, providing a framework for dissecting primary and secondary transcriptional responses. Further, we show that early transcriptional expression of *WRKY70*, within minutes of NHP treatment, mediates secondary transcriptional reprogramming that is required for NHP signaling, ROS production, and SAR.

### Early short-lived NHP transcriptional responses associate with general stress

In the early response to NHP, we identified a primary wave of NHP signaling that was short lived and activated transcriptional responses mirroring wounding/DAMP and JA signaling in a partially SA-dependent manner. Of the pathways activated by NHP, we found a considerable enrichment of known JA-response genes, including *MYC2* and *JAZ10* (Fig. 2C, Table 1, Table 2). Given the established antagonism between JA and SA signaling pathways (Pieterse et al., 2012), it was surprising to find the majority of genes in the early transient cluster were SA-dependent (Fig. 3A). These findings suggest that part of the early NHP transcriptional response depends on synergism between JA signaling components and pre-infection levels of SA. Our data support a model where general stress response signatures are activated by NHP treatment in an early pulse of gene expression that acts synergistically with SA to modulate activation of defense pathways.

Previous work has described a strong, systemic JA response in Arabidopsis leaves infected with an avirulent *Pst* strain after 4 h, suggesting JA signaling pathways are involved in SAR development (Truman et al., 2007). The JA response detected by Truman *et al*. in the systemic leaves of avirulent *Pst*-treated plants after 4 h may be reflective of the early and transient response to NHP observed in our transcriptional analysis. These JA- and early NHP-responsive marker genes may prove useful for determining optimal time points in testing the systemic translocation of NHP. Furthermore, although genetic studies contest the requirement for JA biosynthesis and signaling in SAR (Attaran et al., 2009), it remains a possibility that differences in primary pathogen treatment or secondary pathogen infection may shift the requirement for JA signaling in SAR establishment and warrant further study.

### Early strong NHP responses activate a primary wave of *WRKY* transcription

We discovered another primary wave of early NHP signaling that was partially SA-independent and exhibited strong induction of *WRKY* gene transcription within minutes of NHP treatment, persisting several hours after treatment. All six *WRKY* TF genes in the early stable upregulated gene cluster were activated by NHP treatment in the *sid2-2* mutant, and although some showed delays in the timing of induction in *sid2-2* seedlings, all six were strongly upregulated by 3 h (Table 1, Table 2). These findings are intriguing given that the expression of these *WRKY* TFs is known to be induced by SA and SA-analogs in an NPR1-dependent manner (Kalde et al., 2003; Wang et al., 2006). The ability of NHP to induce an early wave of *WRKY* expression, independent of elevated SA, suggests NHP and not SA is the primary signal driving this early response.

The early and strong induction of *WRKY* transcription, including two sets of homologs belonging to WRKY group III (*WRKY38* and *WRKY62* as well as *WRKY54* and *WRKY70*), suggested WRKY TFs may play a role in driving the secondary transcriptional response to NHP. Members of WRKY group III are known to play both positive and negative regulatory roles in defense responses (Kalde et al., 2003; Pandey and Somssich, 2009). Consistent with this, we found CREs of WRKY TFs to be abundant in the promoters of genes, both up and downregulated, several hours after NHP treatment (Fig. 4C, Supplemental Fig. 5). For example, the CRE bound by WRKY70, as defined by the DAPseq Plant Cistrome Database (O’Malley et al., 2016), was enriched in the promoters of genes upregulated by NHP at 3 and 6 h (Fig. 4C). Other canonical motifs bound by WRKY70, such as the W-box and WT-box (Rushton et al., 2010; Machens et al., 2014), were also found in the promoters of genes upregulated several hours after NHP treatment (Fig. 6A). Taken together, these findings link the early NHP-driven induction of *WRKY* expression to the subsequent up and downregulation of WRKY-targeted genes 3-6 h after initial NHP treatment. Based on these findings, we hypothesize a model of early NHP signaling wherein activation of *WRKY* transcription is a primary response to NHP signaling that is followed by a secondary wave of expression driven through WRKY regulation.

While not the focus of this study, it is noteworthy to mention that TGA TFs are also associated with the primary wave of NHP mediated transcription. CREs of TGAs were abundant in the promoters of early NHP upregulated genes across all time points sampled, particularly those in the early strong cluster (Fig. 4B). These findings are in agreement with recent studies demonstrating TGAs are redundantly required for NHP-induction of SAR and SAR marker genes (Nair et al., 2021; Yildiz et al., 2023). Our data provide further evidence highlighting the importance of TGAs in NHP-driven signaling and suggest that a NPR1/TGA complex, or a key interactor of this complex, directly relays the NHP signal.

### Late weak and late strong genes define the secondary wave of transcription

Finally, our RNA-seq analysis also revealed a secondary wave of NHP signaling defined by late weak and late strong upregulated genes. These genes are associated with pathogen defense, SAR, and SA responses, and showed features of SA-dependent and SA-independent transcriptional induction (Fig. 2C, Fig. 3A). Upregulation of SAR-associated genes by NHP is consistent with previous studies (Bernsdorff et al., 2016; Yildiz et al., 2021; Yildiz et al., 2023). Our study reveals that transcriptional activation of these SAR-associated genes occurs as early as 3-6 h following detection of the NHP signal. Interestingly, 24 genes from the late weak cluster and 4 genes from the late strong cluster are SA-independent within 6 h of NHP treatment but SA-dependent 24 h post NHP (Yildiz et al., 2021). Among this group are key SA and NHP biosynthetic and regulatory genes including *ALD1*, *UGT76B1*, *SARD1* and *NPR3*. Together, these data suggest NHP is sufficient to drive gene expression over a short time frame but requires SA accumulation for sustained expression, thus highlighting the importance of interrogation into early NHP transcriptional changes. Further elucidation of a timeframe for early NHP responses should prove useful in studying other aspects of NHP biology, such as the biosynthesis and translocation of NHP and the identification of NHP sensors and their respective downstream signaling pathways.

### Primary NHP-responsive gene *WRKY70* mediates secondary transcriptional changes required for NHP-elicited SAR, signaling, and ROS production

We found *WRKY70* expression is induced within 15 min of NHP treatment and is required for full NHP-elicited SAR, as NHP did not enhance resistance in distal leaves of the *wrky70-1* mutant and could not achieve the same level of resistance observed in NHP-treated wild type plants (Fig. 5A). WRKY70 is known to repress many pathogen-inducible genes in the absence of infection, but is also required for full activation of pathogen-inducible genes during infection (Zhou et al., 2018). We observed enhanced resistance to *Psm* infection and enhanced transcript abundance of defense genes *SARD1*, *PR1*, *PR2*, and *PR5* after mock treatment (Supplement Fig. S7), consistent with a repressive role for WRKY70. However, NHP treatment did not enhance resistance nor increase transcript abundance of these genes in *wrky70-1* (Fig. 5A, Supplement Fig. S7). Furthermore, expression of the NHP-upregulated early strong gene *BDA1* was completely abolished in *wrky70* mutant lines (Fig. 6B, Supplemental Fig. S8), demonstrating early expression of *WRKY70* is required to drive and modulate a subset of the secondary transcriptional responses of NHP signaling.

Evidence supports a SA- and NPR1-independent defense pathway driven by WRKY70 (Shah et al., 2001; Zhang et al., 2010b). We found that NHP was sufficient to induce expression of *WRKY70* in the *sid2-2* mutant within 30 min of treatment (Table 1), suggesting SA biosynthesis is dispensable for the early transcriptional activation of *WRKY70*. Likewise, WRKY70-regulated genes *BDA1* and *CML10* are induced within 3-6 h of NHP signaling partially independent of SA (Table1). Notably, 24 h after NHP treatment, *BDA1* and *CML10* are also upregulated in both *sid2-1* and *npr1-3* mutants (Yildiz et al., 2021). It is therefore possible that NHP signaling activates a SA- and NPR1-independent defense pathway through the action of WRKY70.

Studies into the MAMP-signaling pathway functioning through the receptor-like protein SNC2 suggest a role for WRKY70 and BDA1 in PTI responses such as ROS production (Yang et al., 2012). While flg22-elicited ROS production was not significantly different between mock treated wild type and *wrky70* mutants, the enhanced production of flg22-elicited ROS following NHP and SA pretreatment was compromised in *wrky70* mutants (Fig. 7, Supplemental Fig. S9). Our findings indicate WRKY70 positively contributes to the NHP and SA signaling pathways that prime PTI. Further, our discovery that NHP primes plants for enhanced production of flg22-elicited ROS is consistent with reports of NHP priming Arabidopsis for enhanced metabolic and transcriptional responses following flg22 treatment (Löwe et al., 2023). Our work links WRKY70-dependent signaling pathways to this NHP-primed PTI response.

## Conclusion

Our work describes the early transcriptional response to NHP and highlights key genes and signaling pathways that may contribute to the physiological processes regulated by this important bioactive molecule. It also provides a foundation for better understanding the transcriptional regulators, particularly the role of WRKY70, and gene networks that define early NHP signal transduction.

## Material and methods

### Plant materials and growth conditions

*Arabidopsis thaliana* (Arabidopsis) lines used in this study were in the Col-0 ecotype (wild type) with mutant lines as follows: *sid2-2* (Wildermuth et al., 2001) and T-DNA insertion lines *wrky70-1* (SALK_025198) and *wrky70-2* (GABI_324D11) (Li et al., 2006), *wrky38* (WiscDsLox489-492C21), *wrky54* (SALK_017254), and *wrky59* (SALK_039436) (Wang et al., 2006), *wrky51* (SALK_022198) (Yan et al., 2018), and *wrky62* (SM_3_38820) (Kim et al., 2008). All mutant lines were obtained from the Arabidopsis Biological Resource Center (ABRC). The presence of T-DNA insertions were confirmed with the genotyping primers listed in Supplemental Table S1. Plants were grown on soil in a controlled growth chamber at 80% humidity and 22°C on a 10 h light/14 h dark cycle for gene expression and bacterial growth assays. For seedlings grown hydroponically, seeds were surface sterilized and stratified at 4°C in darkness for 3 days. Seeds were placed into sterile six-well plates (15-18 seeds per well) with half-strength Murashige and Skoog (MS) medium, 0.5% (w/v) sucrose, 0.05% (w/v) MES at pH 5.7. Plates were covered with a lid, sealed with micropore tape (3M), and placed in a chamber on a 10 h light/14 h dark cycle. Liquid medium was agitated twice daily to facilitate gas exchange and after five days of growth, spent medium was replaced with fresh MS.

### Bacterial strains and growth conditions

*P. syringae* pv. *maculicola* strain ES4326 (*Psm*) and *P. syringae* pv. *tomato* DC3000 (*Pst* DC3000) carrying pVSP61 + *avrRpt2* (*Pst avrRpt2*) were used in this study. Growth conditions for all bacterial strains are as previously described (Holmes et al., 2021).

### Chemical stocks

Two sources of NHP were used in this study, one synthesized by Elizabeth Sattely’s lab as previously described (Chen et al., 2018) and the second obtained from Med Chem Express. Gene expression and NHP-elicited SAR assays showed similar results for both sources of NHP. Pipecolic acid stocks were made from L-pipecolic acid (Oakwood Chemical). Salicylic acid stocks were made from sodium salicylate (Mallinckrodt).

### Chemical treatment of plants for qRT-PCR assays

For gene expression by qRT-PCR of 4.5-week-old, soil-grown Arabidopsis, three leaves were infiltrated with sterile water or NHP dissolved in water at the indicated concentrations. The three treated leaves were collected and flash frozen at the indicated time points. For the time course of gene expression in *wrky70-1 and wrky70-2* mutants, three leaves were treated with sterile water or 1 mM NHP. One leaf from the same plant was sampled at 6, 9, and 12 h and pooled with a second plant of the same condition for one replicate. Samples collected at 0 h were untreated before collecting.

For gene expression in Arabidopsis seedling-based assays, plants were grown as described for seedling hydroponics and treated at 10-days-post-germination. Before addition of treatment, all liquid medium was removed from plate wells and replaced with fresh MS medium, seedlings were allowed to recover under lights for at least 1 h, followed by application of 2x solutions of Pip, NHP, or mock in MS medium. The 15-18 seedlings within a single well were pooled and treated as a single replicate, with three replicates sampled per condition. Following treatment, seedlings were blotted dry, and flash frozen at the indicated time points.

### RNA isolation and qRT-PCR

Total RNA was isolated from leaves and seedlings using TRIsure™ reagent (Meridian Bioscience) according to the manufacturer’s instructions. 5 µg of RNA was used to synthesize cDNA by oligo dT (New England Biolabs) and reverse transcriptase (Thermo Fisher), followed by dilution of cDNA. The qRT-PCR reactions were performed on a MJ Opticon 2 (Bio-Rad) using Green Taq DNA polymerase (GenScript) with EvaGreen Dye (Biotium) for amplification and detection. Three technical replicates were performed per sample and three to four biological replicates were included for each condition. Expression values were normalized to the reference gene *UBC21* (AT5G25760) for relative expression determined using 2^−ΔCt^ and normalized to both *UBC21* and mock treated samples for fold change using 2^−ΔΔCt^. Genes assayed include *FMO1* (AT1G19250), *ICS1*/*SID2* (AT1G74710), *UGT76B1* (AT3G11340), *WRKY38* (AT5G22570), *SARD1* (AT1G73805), *PR1* (AT2G14610), *PR2* (AT3G57260), *PR5* (AT1G75040), *BDA1* (AT5G54610), *PROSCOOP4* (AT5G44568), and *CML10* (AT2G41090). Primers used in these experiments can be found in Supplemental Table S1.

### RNA-seq treatment and library preparation

Seedlings used for mRNA isolation were grown hydroponically as described above. Six-well plates contained equal numbers of Col-0 and *sid2-2* seedlings, with one genotype per plate and 15 seedlings per well. At 10-days-post-germination, all liquid medium was removed from each well and fresh MS medium was added to the seedlings. Seedlings were placed back into the growth chamber for an hour to acclimate, followed by treatment with 2x NHP (final concentration 0.5 mM) or mock (MS medium). Three wells of pooled seedlings were collected for each genotype, treatment, and time (15 min, 30 min, 3 h, and 6 h) combinations, blotted dry, and flash frozen for mRNA isolation.

Total RNA was extracted from pooled seedlings using TRIsure™ reagent (Meridian Bioscience) according to the manufacturer’s instructions. Total RNA was then divided with 5 µg used for cDNA synthesis and marker gene expression to assess quality of treatment, as described for qRT-PCR experiments, and 5 µg was used for cDNA library preparation. The cDNA library was prepared using NEBNext® Ultra™ II RNA Library Prep Kit for Illumina® with NEBNext® Poly(A) mRNA Magnetic Isolation Module and then multiplexed via PCR amplification using NEBNext® Multiplex Oligos for Illumina® according to the manufacturer’s instructions (New England Biolabs). Library concentration was quantified with Qubit™ dsDNA HS Assay Kit (Thermo Fisher Scientific) followed by measurement of quality and average cDNA length using the DNA 1000 Series II chip on a 2100 Bioanalyzer (Agilent Technologies). The multiplexed cDNA libraries were then pooled (two sets of 36 libraries) and sequenced using both lanes of the NovaSeq 6000 (Illumina) S1 flow cell with run type 100 bp paired-end reads at the Genome Sequencing Service Center by the Stanford Center for Genomics and Personalized Medicine, supported by the grant award NIH S10OD025212.

### RNA-seq data analysis

Read data was initially assessed using FastQC. CLC Genomics Workbench version 11.0.2 (QIAGEN) was used for trimming, discarding reads trimmed below 20 bp, and then mapped to the Arabidopsis TAIR10 genome with the auto-detect paired distances option deselected. Differential expression analysis was performed using DESeq2 (Love et al., 2014) with mock treated samples compared to NHP treated samples for each respective time point and genotype. Genes were considered differentially expressed if they returned an adjusted *P-*value below 0.05 (*P*_adj_ < 0.05). The complete list of early NHP-responsive transcripts is provided in Supplemental Table S3 and the raw sequencing data deposited in the Gene Expression Omnibus (GEO) under accession code GSE263276.

### Exploratory analysis of differentially expressed genes

To define patterns of early expression in response to NHP treatment, differentially expressed genes were considered upregulated in wild type if they had a log_2_(FC) > 1 at any time point and downregulated with a log_2_(FC) < −1 at any time point. The sets of up and downregulated genes were then hierarchically clustered by log_2_(FC) over the four timepoints using the hclust function and dendextend packages in R, with Euclidean distances and Ward.D2 linkage (Galili, 2015). For each defined cluster in wild type, the log_2_(FC) values were averaged at each time point to visualize broad trends in expression.

The PANTHER classification system accessed via TAIR was used for gene ontology enrichment analysis (https://www.arabidopsis.org/tools/go_term_enrichment.jsp; (Mi et al., 2021)). Version 17.0 of the PANTHER overrepresentation test was used (Fisher’s exact test with FDR correction) and matched to the GO database 2021_03 release.

### Promoter and *cis*-regulatory element enrichment analysis

Araport11 sequences 1000 bp upstream of the TSS of NHP-responsive genes were downloaded from TAIR (https://www.arabidopsis.org/tools/bulk/sequences/index.jsp) for each time point and genotype grouping of the NHP upregulated and downregulated genes. A less stringent cut-off of log_2_(FC) > 0 (upregulated) or log_2_(FC) < 0 (downregulated) with a *P*_adj_ < 0.05 was used to select for NHP-responsive genes. Motif identification and enrichment of *cis*-regulatory elements (CREs) was performed using SEA, part of the MEME suite (Bailey et al., 2015; Bailey and Grant, 2021) and compared to the published library of TF binding sites found within the DAPseq database (O’Malley et al., 2016). Only CREs from the DAP-seq library were used in this study, omitting CREs from the ampDAP-seq library (where DNA modifications have been removed). Following the workflow detailed by Bjornson et al. (2021), relative enrichment was determined against the same three sets of randomly sampled genes detected in this RNA-seq experiment for all genotypes and time points tested. The identified motif was considered enriched if it returned a significant *P-*value (< 0.05) relative to at least two of the three randomly sampled gene sets. *P-*values for significantly enriched TF-binding motifs were then averaged for visualization.

### Measurement of bacterial growth in plants

Chemical treatment of leaves for NHP-elicited SAR assays were performed as described in (Chen et al., 2018) with minor alterations. At 4.5-weeks-old, three lower leaves of wild type and mutant Arabidopsis plants were infiltrated with sterile water, 0.5 mM NHP or 1 mM NHP. After 24 h, one untreated distal leaf of each plant was inoculated with a 1 x 10^5^ CFU/mL suspension of *Psm*. The distal infected leaf was always separated by one leaf from the original three treated (i.e., primary treatment in leaf no. 7-9 then infected leaf is no. 11). Inoculated plants were covered with a dome to increase humidity and 3 dpi the titer of *Psm* in the distal leaves was quantified. Leaf discs were homogenized in 1 mL of 10 mM MgCl_2_, plated in triplicate in a dilution series on nutrient yeast glycerol medium supplemented with 1.5% wt/vol agar (NYGA) with rifampicin (100 μg/mL), incubated at 28 °C for 1.5 d, and bacterial colonies counted. For each individual experiment, 4-12 plants were tested per condition.

Pathogen priming of leaves for SAR assays were performed as previously described (Chen et al., 2018). Three lower leaves of wild type and mutant Arabidopsis plants (4.5-weeks-old) were infiltrated with 10 mM MgCl_2_ or a 5 x 10^6^ CFU/mL suspension of *Pst avrRpt2* in 10 mM MgCl_2_. After 48 h, one distal untreated leaf of each plant was inoculated with a 1 x 10^5^ CFU/mL suspension of *Psm* and plants were covered with a dome to increase humidity. At 3 dpi, the titer of *Psm* in the upper leaves was quantified as described above for NHP-elicited SAR. For each individual experiment, 8 plants were tested per condition and experiments repeated twice with similar results.

For bacterial growth measurements in unprimed leaves of Arabidopsis wild type and mutant plants, a suspension of 1 x 10^5^ CFU/mL *Psm* was infiltrated into two young leaves per plant. Plants were placed under a dome and 3 dpi the two infected leaves were pooled and bacterial titer quantified for 12 plants per genotype.

### Oxidative burst assay

Four leaf discs (4 mm diameter) from 5-week-old Arabidopsis plants (n = 12) were incubated on water, 500 µM NHP, or 100 µM SA in a 96-well plate (one leaf disc per well) for 24 h. After pretreatment, ROS was measured by addition of flg22 (100 nM) in a solution of 20 mg/mL horseradish peroxidase and 200 µM Luminol (Sigma) (GómezLGómez et al., 1999), followed by immediate measurement of luminescence in a Synergy H1 Microplate Reader (Biotek). Relative luminescence units (RLU) are reported.

### Statistical analyses

Two-tailed Student’s *t* tests were performed to determine statistically significant differences between the different conditions for measurements of bacterial growth using the stat_compare_means function of the ggpubr package (Kassambara, 2023). *P* values < 0.05 were considered statistically significant and all significance levels are indicated in the figure legends (\**P* < 0.05, \*\**P* < 0.01, \*\*\**P* < 0.001, \*\*\*\**P* < 0.0001). Statistical significance of qRT-PCR assays was determined using Mann-Whitney *U* test or one-way ANOVA with post hoc Sidak test using the ggpubr and lsmeans R packages, respectively (Lenth, 2016).

## Acknowledgements

We thank the Mudgett laboratory for feedback on the research and manuscript. We also thank Zhiyong Wang, Dominique Bergmann, Sharon Long, and Virginia Walbot for instrument use and intellectual discussion. This work was supported by National Science Foundation IOS-2026368 (to M.B.M. and E.S.S) and National Science Foundation Graduate Research Fellowship DGE-1656518 (to J.F.).

## Author contributions

J.F., J.-G.K., E.S.S. and M.B.M contributed to the study design; J.F. and J.-G.K. performed research; J.F., J.-G.K., E.S.S. and M.B.M analyzed data; J.F., and M.B.M. wrote the manuscript; J.F., J.-G.K., E.S.S. and M.B.M. reviewed and revised the manuscript for accuracy and final approval.

