## Supplemental Figures for "Transcriptome analysis reveals role for WRKY70 in early *N-*hydroxy-pipecolic acid signaling"

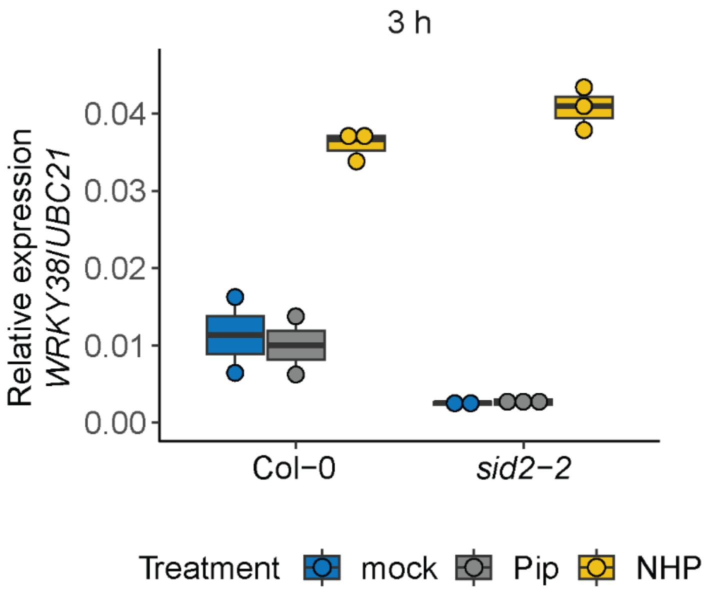


**Supplemental Figure S1** NHP induced *WRKY38* expression is SA-independent. Hydroponically grown wild type (Col-0) and *sid2-2* Arabidopsis seedlings were treated at 10-days-post-germination (dpg) with mock, 1 mM Pip, or 1 mM NHP then collected at 3 h for mRNA isolation. *WRKY38* transcript abundance was measured via qRT-PCR and relative expression ($2^{-\Delta Ct}$) for each condition was determined relative to *UBC21* abundance. One biological replicate consists of 15-18 pooled seedlings (n = 2-3).

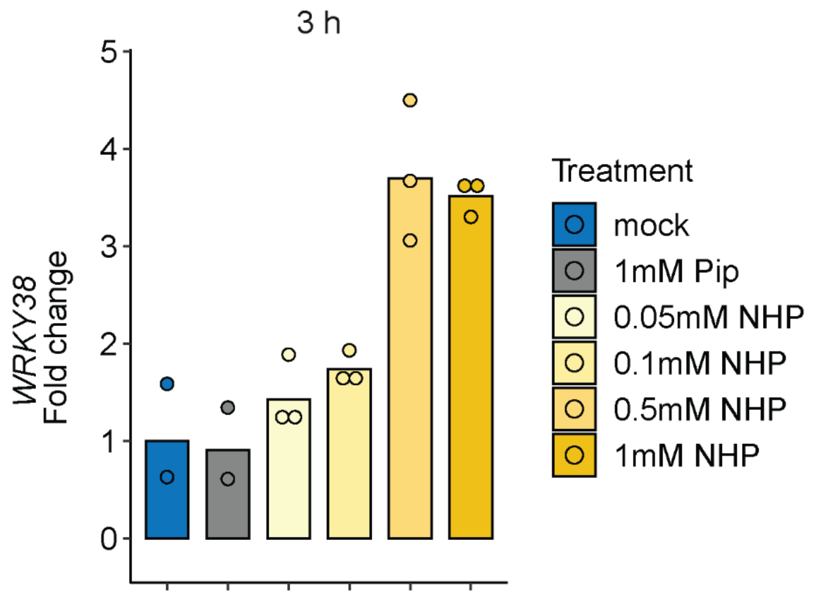


**Supplemental Figure S2** Changes in *WRKY38* mRNA abundance at 3h in response to NHP concentrations.

Hydroponically grown wild type (Col-0) Arabidopsis seedlings were treated with mock, 1 mM Pip, and a range of NHP concentrations (0.05 mM, 0.1 mM, 0.5 mM, and 1 mM) at 10 dpg. After treatment, seedlings were collected at 3 h for mRNA isolation. *WRKY38* transcript abundance was first normalized relative to *UBC21* expression then fold change determined relative to mock ($2^{-\Delta\Delta Ct}$). The data for mock, 1 mM Pip, and 1 mM NHP samples are the same as shown in Supplemental Figure S1. One biological replicate consists of 15-18 pooled seedlings (n = 2-3).

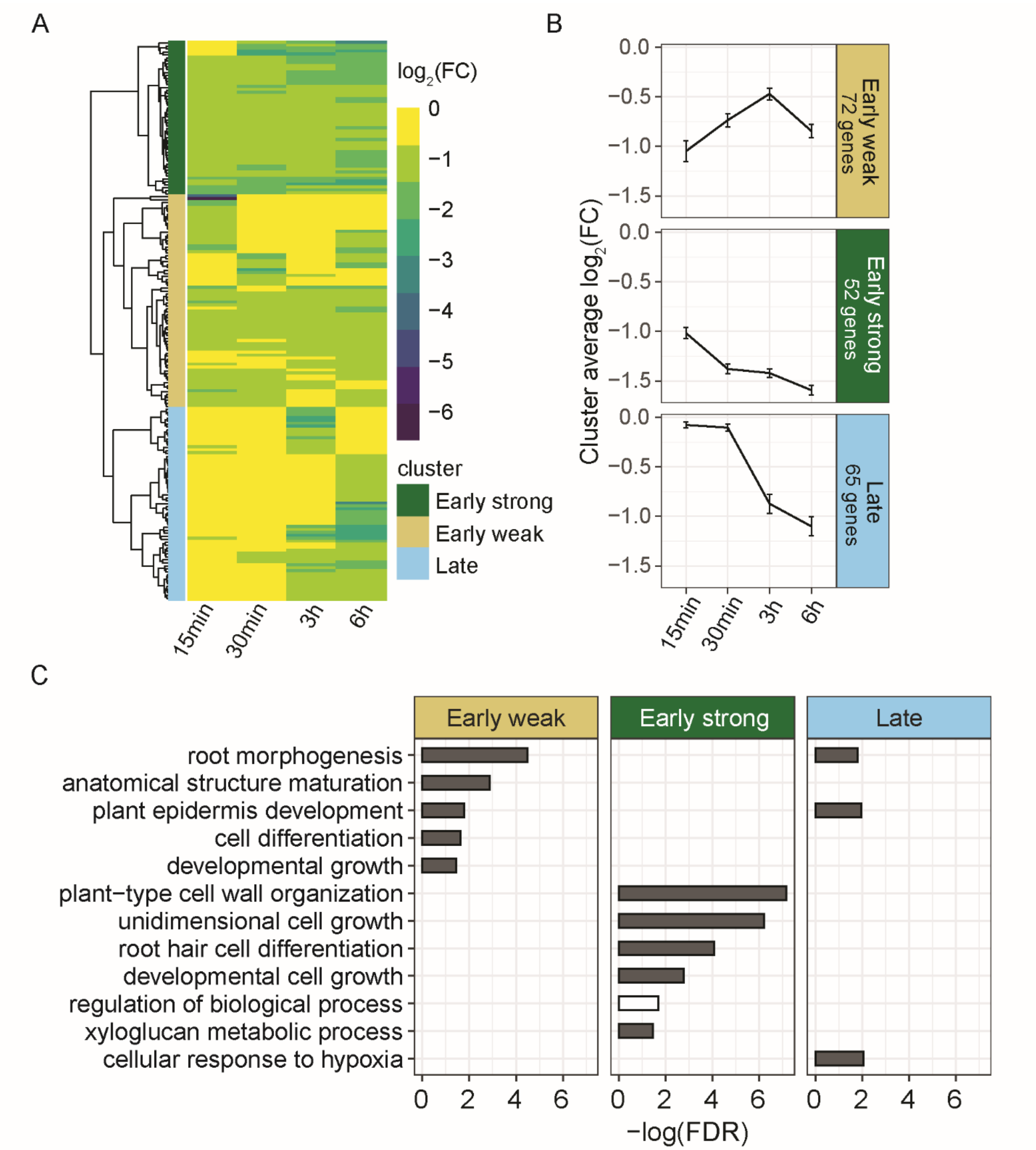


**Supplemental Figure S3** Profile of early NHP-downregulated genes in wild type seedlings.

Three biological replicates of 15 pooled seedlings were performed for each timepoint and genotype. The transcriptome of the treated seedlings was analyzed via RNA-seq and $\log_2$(fold change (FC)) for each time point determined relative to mock treated samples, values were considered significant with a $P_{adj} < 0.05$. A, Heatmap of genes downregulated ($log_2$(FC) < -1, $P_{adj} < 0.05$) in response to NHP treatment in wild type seedlings. Hierarchical clustering was applied to the Euclidean distances determined from the $log_2$(FC) values across the indicated time points, resulting in three distinct clusters of gene expression. B, Average $log_2$(FC) of all genes within each cluster defined in (A) showing the expression trends across the four time points. Error bars represent the ± SEM. C, Biological processes of significantly enriched (gray bar) or depleted (white bar) (FDR < 0.05) Gene Ontology (GO) terms for the genes within each cluster are shown. Bars represent the negative log(FDR) of each significantly enriched GO term. Bars were omitted if the term was not significantly enriched or depleted for the defined cluster.

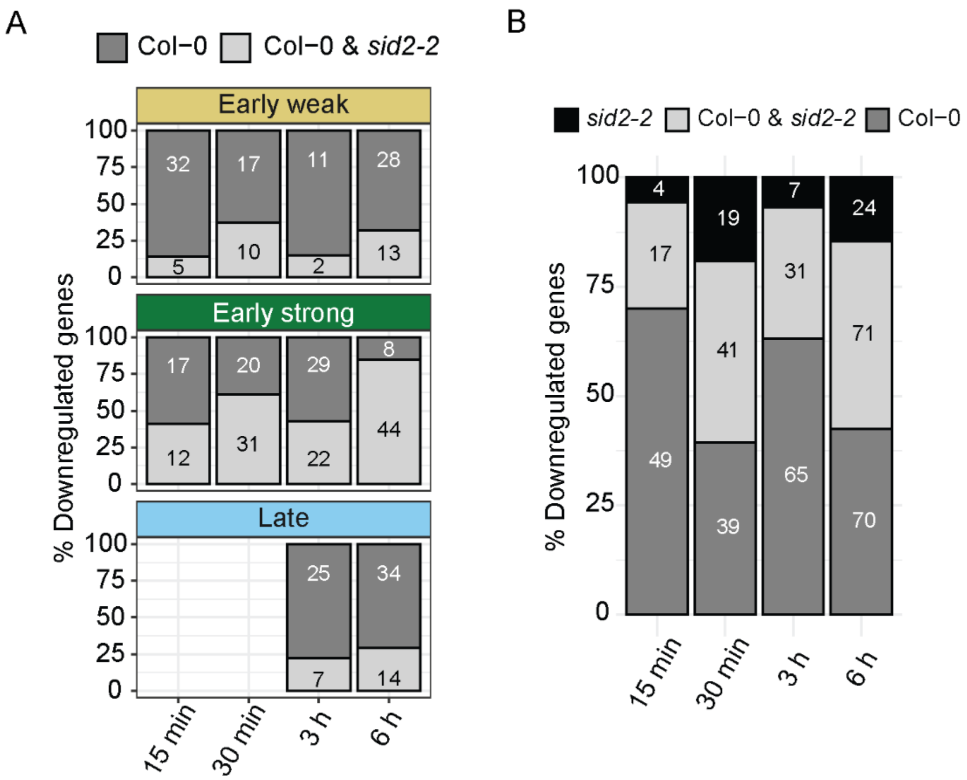


**Supplemental Figure S4** Comparison of NHP downregulated genes in wild type and *sid2-2*.

A, Percent of NHP downregulated genes expressed in wild type (Col-0, SA-dependent) and in both wild type and the *sid2-2* mutant (SA-independent) for the wild type expression clusters defined in Supplemental Figure S3. Total number of genes in each condition is indicated. B, Total number of all NHP downregulated genes ($log_2(FC) < -1$, $P_{adj} < 0.05$) per time point unique to wild type (Col-0), shared between wild type and *sid2-2*, and unique to *sid2-2*. The number of downregulated genes in each grouping is indicated.

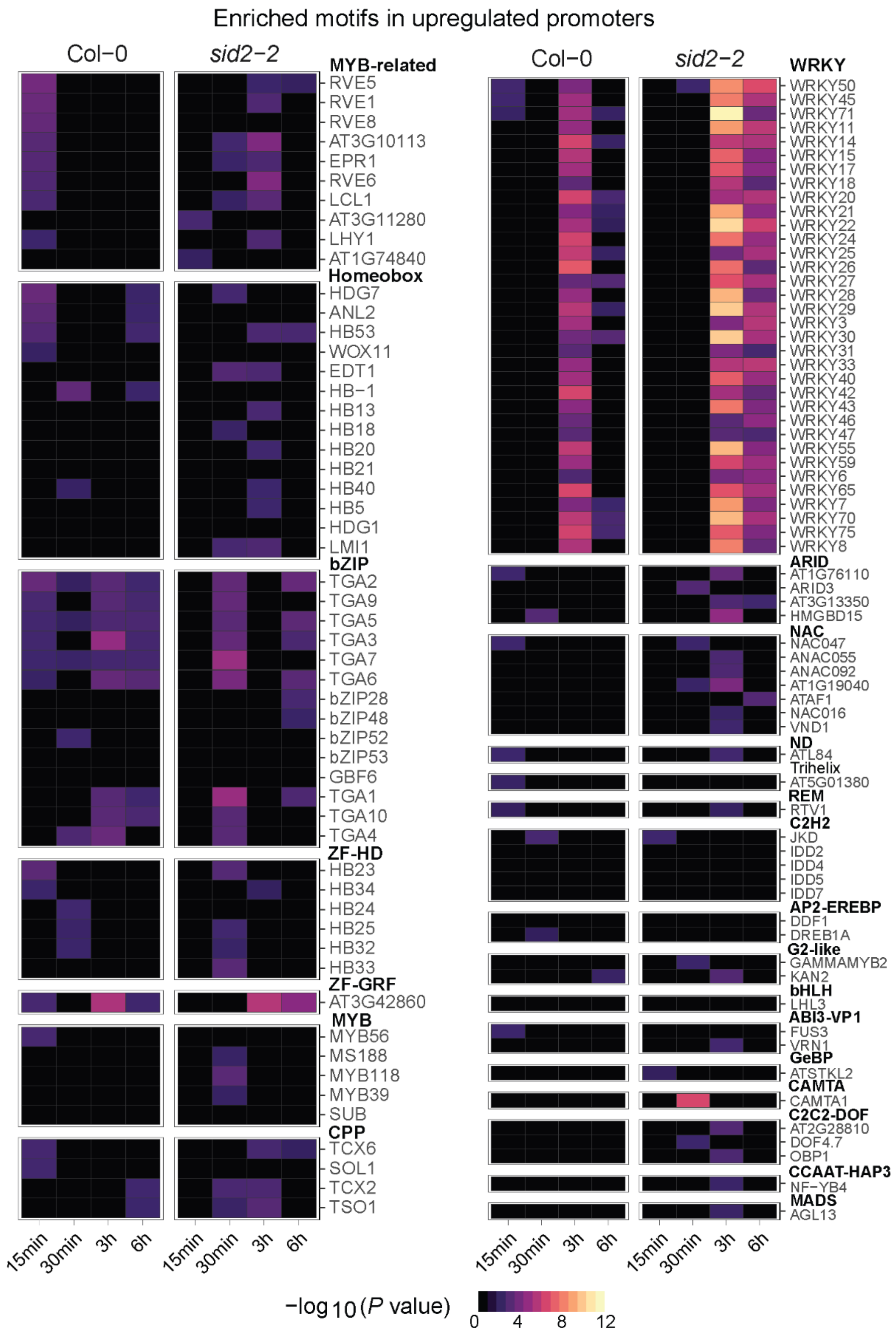


**Supplemental Figure S5** All *cis*-regulatory elements (CREs) in the promoters of NHP-upregulated genes in wild type (Col-0) and *sid2-2* seedlings.

All genes with increased transcript abundance upon NHP treatment ($\log_2$(FC) > 0 and $P_{adj} < 0.05$) were grouped by time point for the indicated genotype. The promoters (-1000 bp upstream of the transcriptional start site) of each grouping were analyzed for the presence of TF-binding motifs (CREs) from the DAPseq database (O'Malley et al., 2016). Enrichment of these CREs was determined relative to TF-binding motifs in promoters pulled from a random sampling of genes detected in this RNA-seq experiment. The -$\log_2$($P$ value) of the enrichment analysis is indicated by the scale bar with black indicating no significant enrichment. The TF family name is indicated in bold above each group and the TF name of the identified binding motif is listed underneath.

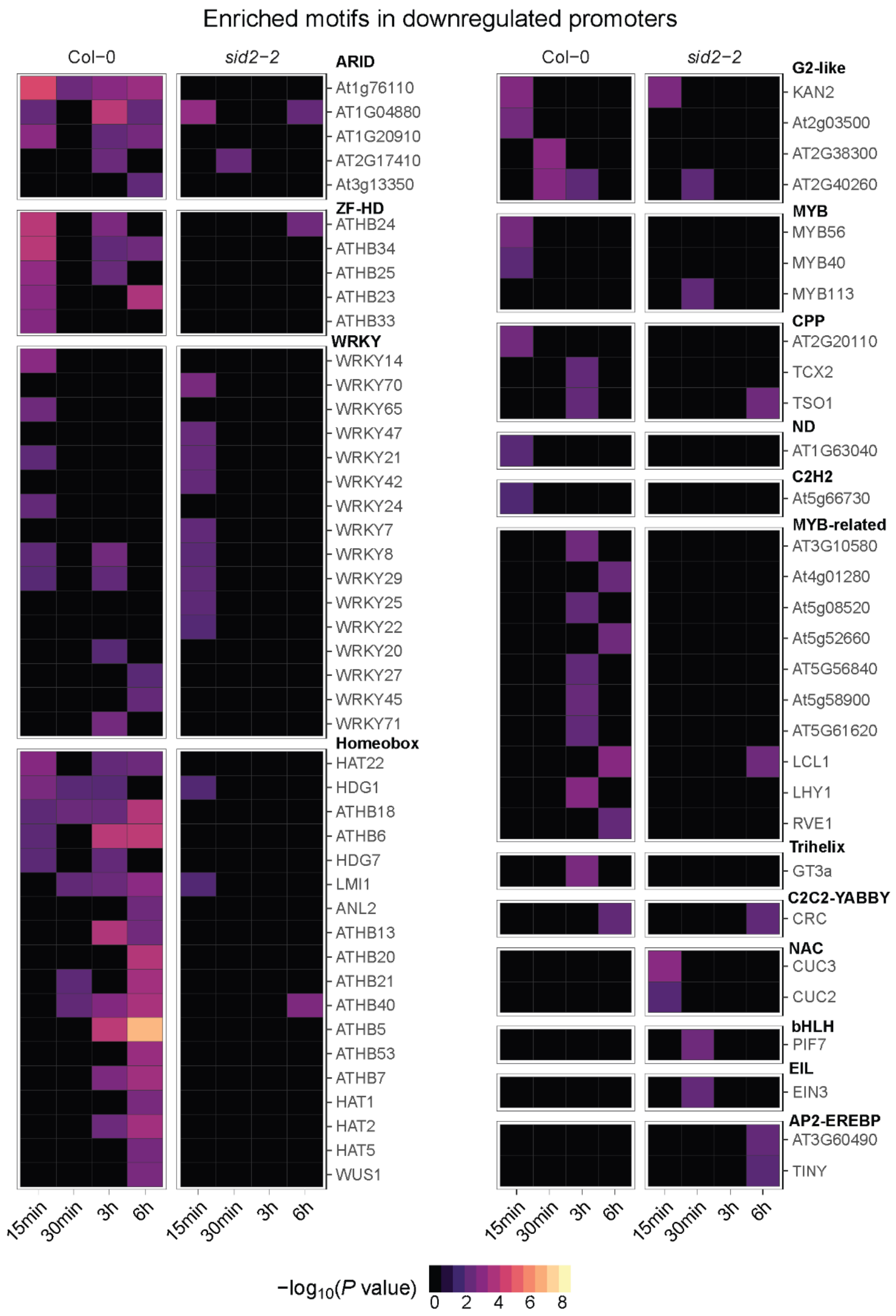


**Supplemental Figure S6** All *cis*-regulatory elements (CREs) in the promoters of NHP-downregulated genes in wild type (Col-0) and *sid2-2* seedlings.

All genes with decreased transcript abundance upon NHP treatment ($log_2$(FC) < 0 and $P_{adj}$ < 0.05) were grouped by time point for the indicated genotype. The promoters (-1000 bp upstream of the transcriptional start site) of each grouping were analyzed for TF-binding motifs (CREs) from the DAPseq database (O'Malley et al., 2016). Enrichment of these CREs was determined relative to TF-binding motifs of promoters pulled from a random sampling of genes detected in this RNA-seq experiment. The -$log_2$($P$ value) of the enrichment analysis is indicated by the scale bar with black indicating no significant enrichment. The TF family name is indicated in bold above each group and the TF name of the identified binding motif is listed underneath.

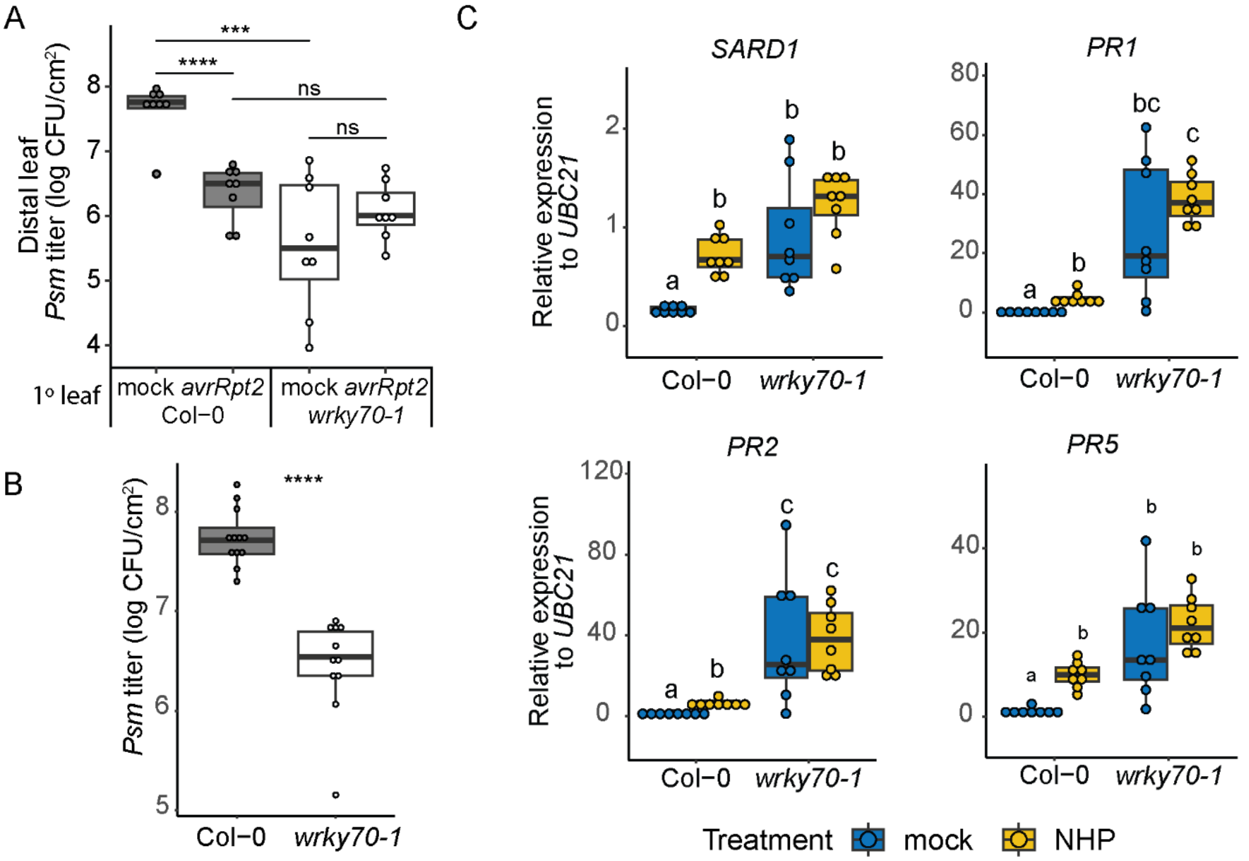


**Supplemental Figure S7** Loss of *WRKY70* results in enhanced resistance to *Psm* infection and elevated levels of defense transcripts independent of NHP treatment.

A, Bacterial growth in the distal leaves of mock and avirulent pathogen treated wild type and *wrky70-1* mutant plants. Three lower (1°) leaves of 4.5-week-old plants were infiltrated with 10 mM $MgCl_2$ (mock) or 5 × $10^6$ CFU/mL of avirulent *Pst avrRpt2*, one distal leaf was inoculated 2 d later with *Psm*, followed by quantification of *Psm* growth 3 dpi (n = 8). B, Resistance to *Psm* in wild type (Col-0) and *wrky70-1* plants without priming treatment. Two leaves of wild type and *wrky70-1* mutant plants were inoculated with a 1 x $10^5$ CFU/ mL suspension of *Psm*, 3 dpi the two leaves were pooled, and bacterial titer measured (n = 11-12). Asterisks indicate significant differences in bacterial titer for A-C (two-tailed t-test; **P* < 0.05, ***P* < 0.01, ****P* < 0.001, *****P* < 0.0001, ns = not significant). D, Expression of defense marker genes *SARD1*, *PR1*, *PR2*, and *PR5*. Three lower leaves of 4.5-week-old plants were infiltrated with water or 1 mM NHP. Treated leaves were then collected at 24 h for mRNA isolation and transcript abundance was determined relative to *UBC21* for each condition ($2^{-\Delta Ct}$). Data is pooled from two separate experiments. Statistical analysis was performed using a one-way ANOVA and post hoc Sidak test, different letters indicated statistical differences between means with $P < 0.05$, n = 8.

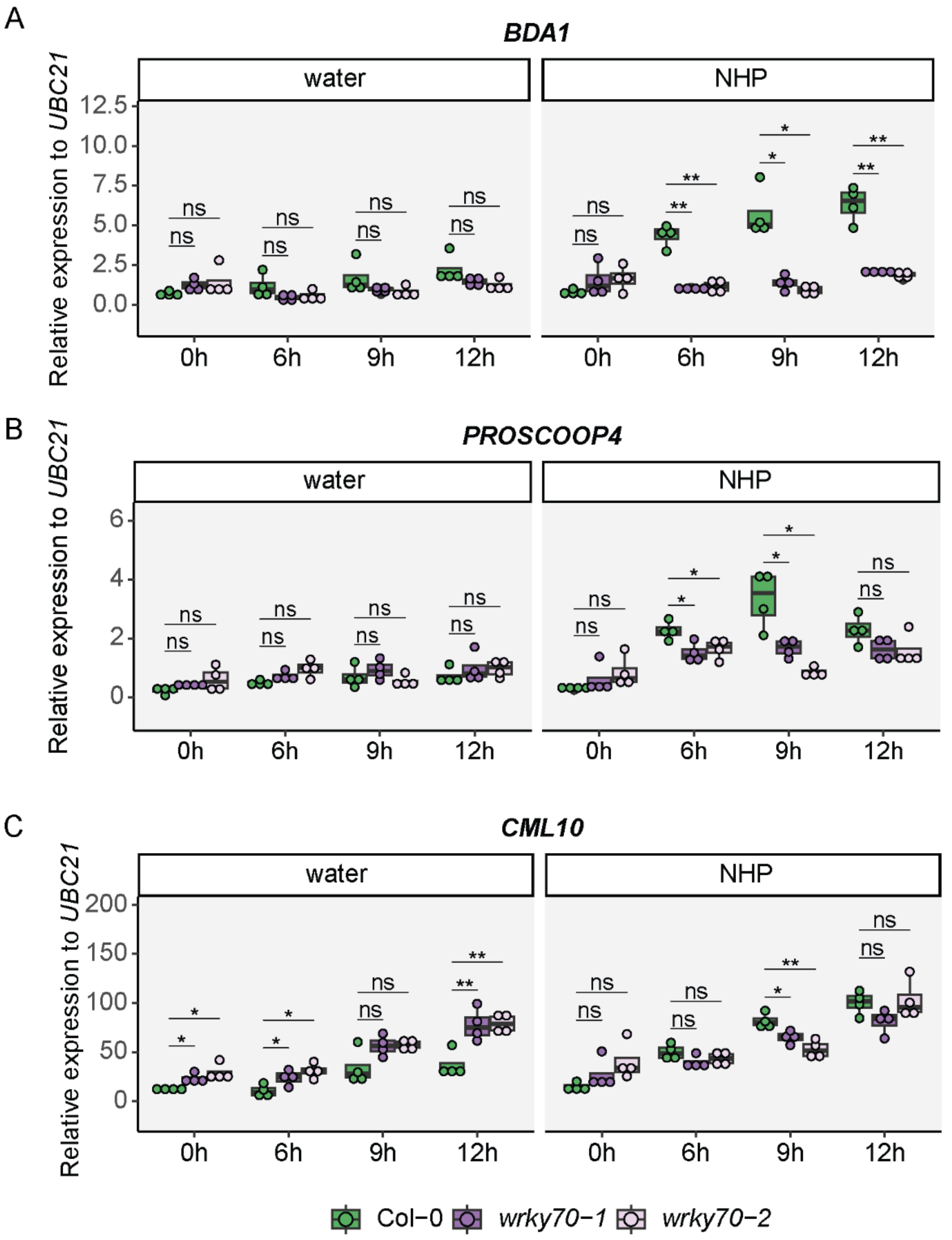


**Supplemental Figure S8** Expression of NHP-responsive transcripts in *wrky70-1* and *wrky70-2* plants.

Expression of *BDA1* (A), *PROSCOOP4* (B), and *CML10* (C) in *wrky70* mutant lines. Three leaves of 4.5-week-old wild type (Col-0), *wrky70-1*, and *wrky70*-2 mutant plants were infiltrated with water or 1 mM NHP. Samples collected at 0 h were untreated before collection. Transcript abundance was determined relative to *UBC21* for each condition ($2^{-\Delta Ct}$). Asterisks indicate a significant difference between wild type and *wrky70-1* or *wrky70-2* at each time point (two-tailed t-test; *$P < 0.05$, **$P < 0.01$, ns = not significant). The data for wild type and *wrky70-1* are the same as shown in Figure 6.

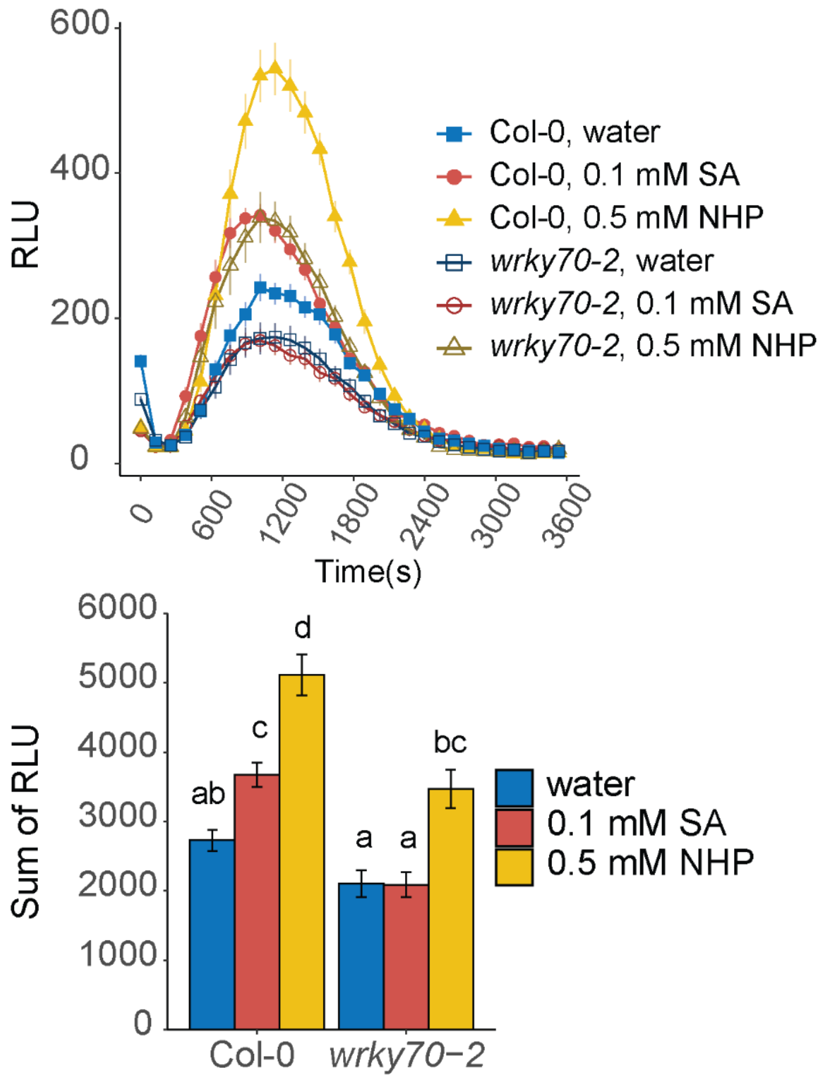


**Supplemental Figure S9** NHP enhanced ROS production is compromised in *wrky70-2* Flg22-elicited ROS quantification in *wrky70-2* plants. Four leaf discs from 5 to 6-week-old wild type and *wrky70-2* plants were pretreated by floating leaf discs on water, 0.1 mM SA, or 0.5 mM NHP for 24 h before treating with 100 nM flg22 in horseradish peroxidase and Luminol. Top, traces of the average relative luminescence units (RLU) over the indicated run time. Each point represents the average of 6 plants (n = 24) from two experiments. Bottom, total quantification of RLUs from top plot. For each leaf disc, RLUs from each time point were summed by condition over the run time and averaged with error bars representing the standard error of the mean. Statistical analysis was performed using a one-way ANOVA and post hoc Sidak test, different letters indicated statistical differences between means with $P < 0.05$.

**Supplemental Table S1**. Primers used in this study

| Description | Primer Name | Sequence (5' - 3') | Reference |
|---|---|---|---|
| **qRT-PCR analysis** | | | |
| *UBC21* | qFW | TCAAATGGACCGCTCTTATC | Sigma-Aldrich |
| | qRV | CACAGACTGAAGCGTCCAAG | |
| *FMO1* | qFW | TCTTCTGCGTGCCGTAGTTTC | Návarová et al., 2012 |
| | qRV | CGCCATTTGACAAGAAGCATAG | |
| *ICS1* | qFW | GCAAGAGTGCAACATCTATATTCTC | Bernsdorff et al., 2016 |
| | qRV | CACAAACAGCTGGAGTTGGA | |
| *UGT76B1* | qFW | TCCCGAGGATTGTTCTCCGAAC | this study |
| | qRV | AACCGGTGAGTCTGCCTTAGTC | |
| *WRKY38* | qFW | GCCCCTCCAAGAAAAGAAAG | this study |
| | qRV | CCTCCAAAGATACCCGTCGT | |
| *SARD1* | qFW | CCTCAACCAGCCCTACGTTA | Truman et al., 2013 |
| | qRV | TAGTGGCTCGCAGCATATTG | |
| *PR1* | qFW | GTGCTCTTGTTCTTCCCTCG | Návarová et al., 2012 |
| | qRV | GCCTGGTTGTGAACCCTTAG | |
| *PR2* | qFW | TCAAGGAAGGTTCAGGGATG | Chen et al., 2018 |
| | qRV | TCGAGATTTGCGTCGAATAG | |
| *PR5* | qFW | AAGGTCATGGATCAGAACAA | Chen et al., 2018 |
| | qRV | GCAGAAAGTGATTTCGTAGT | |
| *BDA1* | qFW | ACGAGCCAGAGGATCTCACATG | this study |
| | qRV | ACTCCTGTAACGTGCAATTCGC | |
| *PROSCOOP4* | qFW | CAGGCCAAACGTCTCCAACAAA | this study |
| | qRV | GCGATAGGAGGTTTGACGATGC | |
| *CML10* | qFW | CTCCGGTGGAAGCAAACTGATG | this study |
| | qRV | GCGCTGCGATGATATTTTTGGC | |
| **Genotype analysis** | | | |
| SALK T-DNA border | LBb1.3 | ATTTTGCCGATTTCGGAAC | |
| SALK_017254 | LP | AACCAGTACCACATCGATGAGAC | |
| | RP | GCTGGTGTTGTTCTCTTGCTC | |
| SALK_025198 | LP | TGATCTTCGGAATCCATGAAG | |
| | RP | CAAACCACACCAAGAGGAAAG | |
| SALK_022198 | LP | TGAATATCTCTCAAAACCCTAGCC | |
| | RP | TTGCTTTCAAACCATGCTTTG | |
| SALK_039436 | LP | GCACGAAAGTAGCCATGAAAG | |
| | RP | TGATTTCTGCAATGAATAAATGTG | |
| GABI kat T-DNA border | 8474_DR35 | ATAATAACGCTGCGGACATCTACATTTT | |
| GABI_324D11 | LP | CAAACCACACCAAGAGGAAAG | |
| | RP | ATGACAAGTCATTCTCCGTGG | |
| pDs-Lox-LT6 T-DNA border | LT6 | AATAGCCTTTACTTGAGTTGGCGTAAAAG | |
| WiscDsLox489-492C21 | LP | ATTTGGTAAACCCAAATTGGC | |
| | RP | CGATGAAGGAGGATAAGAGCC | |
| dSpm T-DNA border | Spm32 | TACGAATAAGAGCGTCCATTTTAGAGTGA | |
| SM_3_38820 | LP | TTACGCGACGACATCAAATAG | |
| | RP | CATAAGCTTGATGAAGCTGGC | |
| *sid2-2* | FW | TGTCTGCAGTGAAGCTTTGG | |
| | RV | CGAAGAAATGAAGAGCTTGGA | |

**Supplemental Table S1** Primer sequences used in this study.

**Supplemental Table S2** Setup of early NHP RNA-sequencing experiment.

**Supplemental Table S3** Full RNA-seq dataset of the early transcriptional response to exogenous NHP in Col-0 and *sid2-2*.

**Supplemental Table S4** Full set of $\log_2$ fold change values grouped by upregulated and downregulated wild type gene clusters.
